# Deciphering the conformations and dynamics of FG-nucleoporins *in situ*

**DOI:** 10.1101/2022.07.07.499201

**Authors:** M. Yu, M. Heidari, S. Mikhaleva, P.S. Tan, S. Mingu, H. Ruan, C.D. Reinkermeier, A. Obarska-Kosinska, M. Siggel, M. Beck, G. Hummer, E.A. Lemke

## Abstract

The ∼120 MDa nuclear pore complex (NPC) acts as a gatekeeper for the molecular traffic between the nucleus and the cytosol. Small cargo readily passes through the transport channel, yet large cargo requires specialized nuclear transport receptors. While the scaffold structure that anchors the NPC in the double-layered nuclear envelope has been resolved to remarkable details, the spatial organization of intrinsically disordered nucleoporins (NUPs) within the central channel remains enigmatic. These so-called FG-NUPs account for about one-third of the total mass of the NPC and form the actual transport barrier. Here we combined site-specific fluorescent labeling in non-fixed cells and fluorescent lifetime imaging microscopy (FLIM) to directly decipher the conformations of an essential constituent of the permeability barrier, NUP98, inside the functioning NPCs using Fluorescence resonance energy transfer (FRET). With detailed measurements of the distance distribution of eighteen NUP98 segments combined with coarse-grained modeling, we mapped the uncharted biochemical environment inside the nanosized transport channel. We found that ‘good-solvent’ conditions for a polymer dominate the inside of the nanosized NPC, expand the FG-domain *in situ* and facilitate nuclear transport, in sharp contrast to the collapsed NUP98 FG-chain in aqueous solution. The combination of fluorescence microscopy, high-resolution electron tomography, and molecular simulation opens a window into the so-far unresolved organization of the FG-NUPs at the center of NPC function, allowing us to reconcile scientific models of nuclear transport.

## Introduction

Intrinsically disordered proteins (IDPs) are flexible, dynamic macromolecules that lack a fixed tertiary structure and can adopt a range of conformations to perform various functions across the cell. IDPs are highly relevant for human physiology and have for example central roles in neurodegenerative diseases and cancer. Intriguingly, IDPs are also key players in phase separation and are involved in the formation of biomolecular condensates (*1*–*9*). In the nanosized nuclear pore complex (NPC) which has a total molecular weight of ∼120 MDa in mammalians, there are hundreds of IDPs enriched in phenylalanine-glycine-residues, called FG-nucleoporins (FG-NUPs) (*10*). The FG-NUPs form a permeability barrier in the central channel of the NPC, which regulates the nucleocytoplasmic transport by restricting the crossing of large cargoes unless they present a nuclear localization sequence or nuclear export sequence (NLS/NES) (*11, 12*). Nuclear transport receptors can specifically recognize the sequence and efficiently shuttle the cargo through the barrier. With recent advances in cryo-electron tomography, crystallography, proteomics, and artificial intelligence (AI)-based structure prediction, ∼70 MDa of the NPC scaffold enclosing the central channel has been resolved with near-atomic resolution (*13*–*20*). However, those structural biology techniques rely on averaging to enhance signals across many individual NPCs, where the signals from the highly dynamic FG-NUPs average out, and therefore the actual transport machinery inside the central channel, i.e., another ∼50 MDa, is not captured leaving a ∼60 nm hole in the center of the scaffold structure. Consequently, the protein conformational state inside the NPC remains elusive and this has led to several partially conflicting hypotheses qualitatively describing the morphologies of the FG-domains in their functional state (*21*–*26*). With ∼30% of the entire eukaryotic proteome being intrinsically disordered, the problem that the conformational state is not easily studied in cells extends far beyond NPC biology. Single-molecule fluorescence tools of purified and labeled proteins have become powerful to probe the conformations of proteins in solution, and in advanced studies it was even shown that it is possible to probe this when microinjected into the cell (*27*–*29*). However, the NPC is only assembled in late mitosis and during nuclear growth in interphase (*30*), and thus requires genetically encoded probes for labeling. The use of popular fluorescent protein-based technologies, however, does not easily enable to extract multiple distance distributions of the same protein, because of the sheer size of the fluorescent labels and the inherently limited labeling freedom.

In this work, we developed a method to quantitatively probe distance distributions of FG-NUPs inside the functional NPCs in non-fixed cells by combining fluorescence lifetime imaging of Fluorescence resonance energy transfer (FLIM-FRET) with a novel site-specific synthetic biology approach. We focused on NUP98 because it is the essential constituent of the NPC permeability barrier and is accessible to this technology (*31*–*33*). By measuring the distance distribution for eighteen labeled chain segments of NUP98 in the NPC using FLIM-FRET, we showed that the FG-domain is exposed to “good-solvent” conditions inside the NPC in the terminology of the Flory polymer model (*34*). This enables the protein to adopt much more expanded conformations in the functional state than in the highly collapsed single chains in solution consistent with “poor-solvent” conditions. With coarse-grained molecular dynamics (MD), we integrated the distance distribution data and the recently solved scaffold structure (*16*) into a molecularly detailed picture of FG-NUP distribution and motion in the central channel of a functional NPC.

## Results

### In-cell site-specific labeling of NUP98 FG-domain in the functional state

Fluorescence measurements of the conformation of FG-NUPs in their functional state with high precision require labeling tags with minimal linkage error and minimal disruption of the structures and functions of the labeled proteins. To this end, we performed site-specific labeling of a non-canonical amino acid (ncAA) *via* genetic code expansion (GCE) (*35*). Thereby, the pyrrolysine orthogonal tRNA/synthetase suppressor pair reassigns the Amber stop codon (TAG), to incorporate the ncAA trans-cyclooct-2-en-L-lysine (TCO*A) at that site. The chemical functionality of this ncAA is then labeled with a clickable organic fluorophore presenting a tetrazine moiety. Thus, the dye gets stably attached to the protein *via* a small chemical linker, leaving minimal disruption to the protein structure and functions. However, one downside of this technique is that it is not mRNA-specific, leading to potential background labeling of untargeted proteins with their naturally occurring stop codons being suppressed. To tackle this problem, we utilized our recently developed synthetic orthogonally translating (OT) film-like organelles to form a distinct protein translational machinery on the outer mitochondrial membrane surface (*36*). These organelles exclusively reassigned two Amber codons for the target FG-NUP and incorporated TCO*A at the two specified sites with high selectivity, ensuring minimal interference for endogenous protein translation and negligible background staining (Fig. 1A and Fig. S1). The incorporated ncAAs were labeled with a mixture of donor and acceptor dyes at a molar ratio of 1:2 (see Methods).

**Fig. 1.**
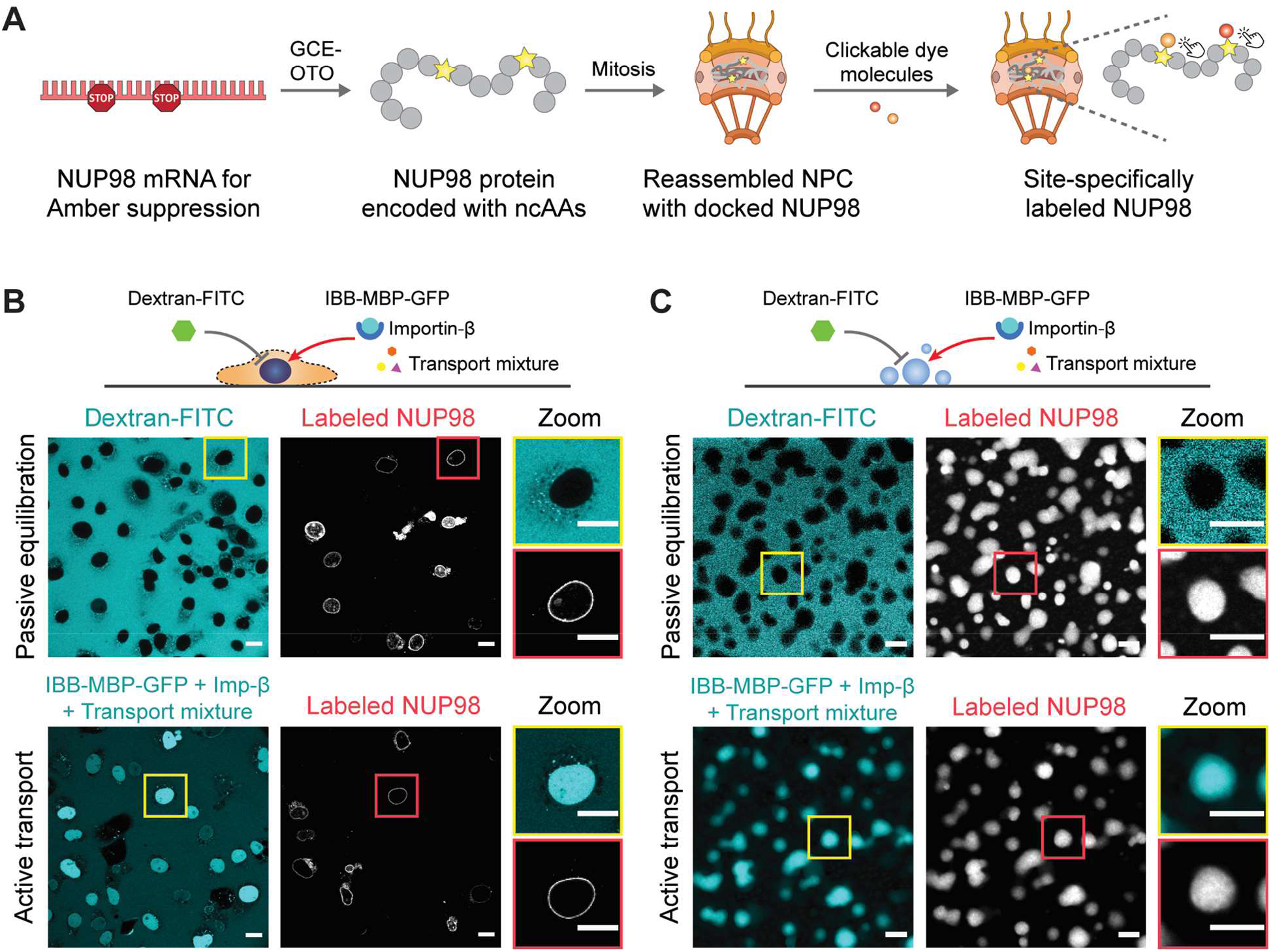
Site-specific labeling of NUP98 in the functional state inside the NPC and comparison with phase-separated condensates *in vitro*. **(A)** Schematic of site-specific labeling of target NUP98 inside the NPC. The genetic code was exclusively expanded for target NUP98. Non-canonical amino acids (ncAAs) were introduced into NUP98 at specific sites by synthetic orthogonal translating organelles (OTO). NUP98 protein encoded with ncAAs docked on the central channel of reassembled NPC during mitosis. Tetrazine-modified dye molecules were added to the cell and reacted with ncAAs *via* click chemistry. **(B)** Passive exclusion assay (70 kDa dextran was excluded) and facilitated/active transport assay (IBB-MBP-GFP supplied with transport mixture was imported) showing that the NPCs were functional with site-specifically labeled NUP98 in COS-7 cells. Scale bar 20 μm. **(C)** Purified NUP98 FG-domain phase-separated *in vitro*. Formed droplet-like condensates recapitulated the permeability barrier-like functions as shown in passive exclusion assay and facilitated transport assay. Scale bar 5 μm.

To verify that our site-specific labeling procedure did not perturb the functionality of the NPC, we performed a permeability barrier functional transport assay on labeled COS-7 cells (Fig. 1B, and Fig. S4A) (*37*). The large inert cargos (70 kDa-Dextran labeled with FITC) were excluded from the nucleus, suggesting that both the permeability barrier and nuclear envelope were intact, whereas IBB-MBP-GFP (a triple fusion of the Importin-β binding domain with maltose-binding protein and green fluorescent protein) supplied with transport mixtures (which contain the nuclear transport receptor Importin-β, see Methods) were actively imported into the nucleus, demonstrating that the NPCs with labeled NUP98 were fully functional.

### FLIM-FRET measurements of NUP98 FG-domain in the NPC

FLIM-FRET is exquisitely sensitive to the spatial distance between donor-acceptor fluorescent dye pairs, independent of fluorophore concentration or excitation intensity, providing an ideal tool to probe the dimension of FG-NUPs in the cellular milieu (*38*). By quantifying donor fluorescence lifetime decrease when it undergoes FRET with an acceptor molecule, we determined the spatial distances between the labeled dye pairs. Here we chose a FRET dye pair of AZDye594-tetrazine and LD655-tetrazine due to their exceptional photostability, spectra far away from cellular autofluorescence, and a suitable Förster radius (R_0_ ∼ 7.7 nm, see Methods).

Given the complexity of the milieu in the central channel of the NPC, the fluorophore properties could vary due to different local environments. By labeling different positions of NUP98 FG-domain with only the donor dye, we showed that the donor lifetimes were practically the same among different labeling positions (see Fig. S5), proving that the fluorophores were unaffected by the cellular environment. Another concern may arise for inter-molecular FRET, i.e., FRET between different NUP98 molecules in the same NPC, reinforced by the crowded central channel. Considering that the endogenous population of NUP98 is substantially more abundant than the ectopically expressed one this, however, seemed unlikely. Nevertheless, to ensure that the mutant NUP98 was not highly overexpressed and the endogenous population of NUP98 remained substantially higher than the exogenous one, we selected the nuclear envelopes with the acceptor intensity per pixel (excited by 660 nm laser) below a determined threshold level before the FRET measurements (Fig. S6, see Methods). Furthermore, to verify that no inter-molecular FRET was measured, we transfected COS-7 cells with a plasmid encoding NUP98 with only a single-Amber-codon (NUP98^A221TAG^) and labeled it with a mixture of donor and acceptor dyes, such that each modified copy of NUP98 could only be labeled with either a single donor or single acceptor dye (Fig. 2B). We measured the average fluorescence lifetime of the donor dye before and after acceptor photobleaching (*39*). In acceptor photobleaching, the acceptor is bleached selectively by a high-power laser. In the presence of FRET, the donor intensity and the fluorescence lifetime will increase after acceptor bleaching. We found the donor signal to be unchanged, thus excluding the existence of inter-molecular FRET (Fig. 2D). Next, we performed the same assay on cells transfected with double-Amber-mutated NUP98 (NUP98^A221TAG-A312TAG^, Fig. 2C). We observed an increase in both the donor intensity on the nuclear rim and its average fluorescence lifetime when the acceptor was photobleached, confirming the occurrence of FRET (Fig. 2E). With these results we validated the experimental setup to be sufficiently sensitive for measuring FLIM-FRET between two labeled sites of NUP98 in fully functional cells.

**Fig. 2.**
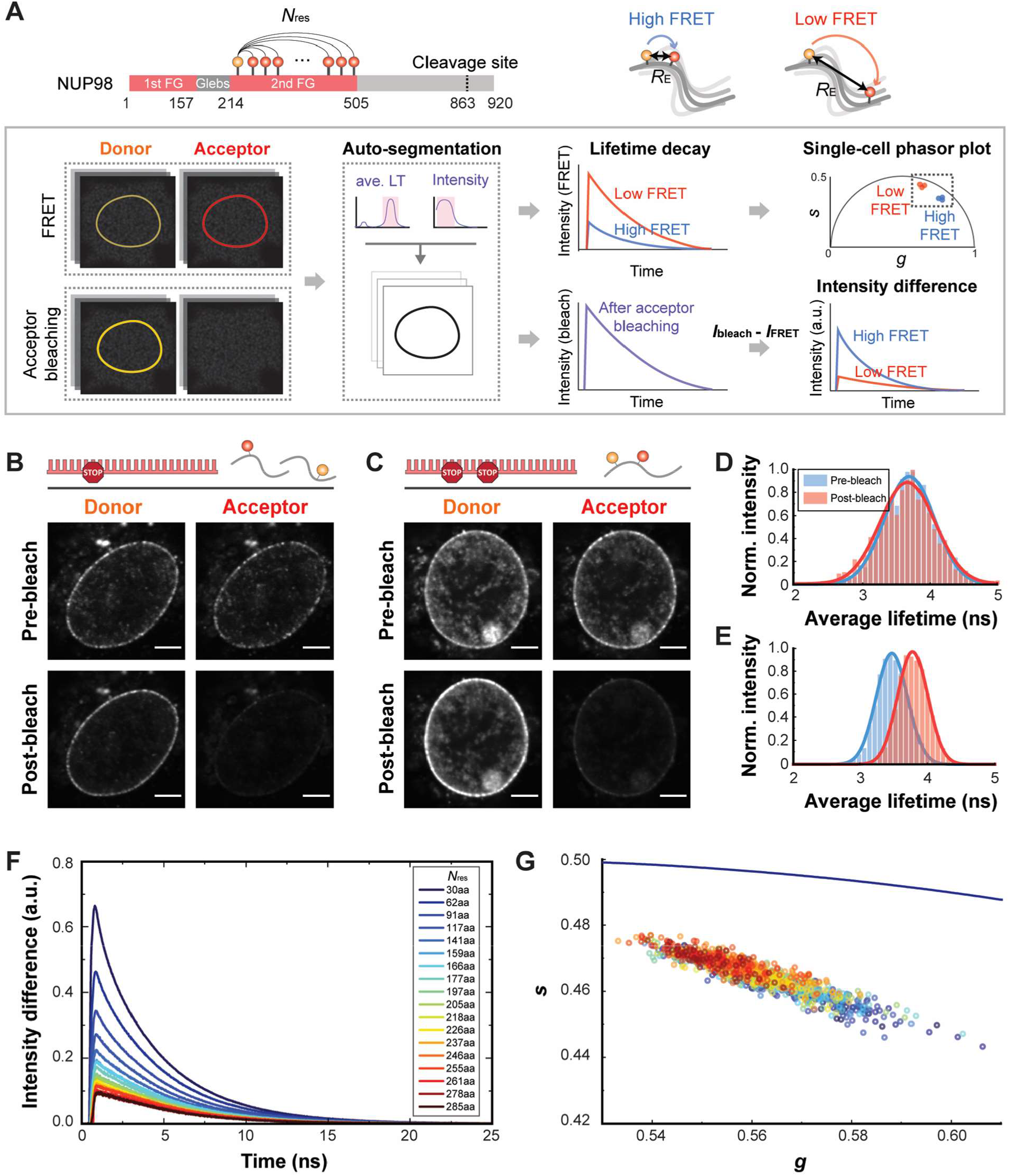
FLIM-FRET measurements of NUP98 FG-domain inside the NPC. **(A)** Schematics of FLIM-FRET analysis pipeline. Different chain segments of NUP98 FG-domain were labeled with a FRET dye pair and the donor fluorescence intensity was measured on a cell-by-cell basis. For each cell, the nuclear rim was selected as a region of interest *via* auto-segmentation, and the measured donor fluorescence intensity profiles before and after acceptor photobleaching were extracted. The fluorescent lifetimes were further converted to a phasor plot *via* Fourier transform for cell-by-cell comparison. **(B)** For a single-Amber-mutated sample labeled with a donor and acceptor dye mixture, the average fluorescence lifetime of the donor dye did not change before and after acceptor photobleaching as shown in **(D)**, indicating that no inter-molecular FRET could be detected. Scale bar 5 μm. **(C)** For a double-Amber-mutated sample labeled with donor and acceptor dye mixture, the average fluorescence lifetime of the donor dye changed before and after acceptor photobleaching as shown in **(E)**, indicating that intra-molecular FRET was detected. Scale bar 5 μm. **(F)** The donor fluorescence intensity differences between before and after acceptor photobleaching for the eighteen chain segments of NUP98 FG-domain. Each profile represents an averaged result of ∼100 cells. The higher peak shows a bigger difference in the intensity profiles, indicating higher FRET efficiency and smaller end-to-end distance. **(G)** Phasor plot showing donor lifetimes of the measured eighteen chain segments on a single-cell basis. The right-shifted dots represent higher lifetime and lower FRET efficiency.

Next, we created a series of NUP98 mutants to form a set of chain segments of different lengths for FLIM-FRET measurements. We chose A221TAG as the reference site and kept it constant while varying the second site along the FG-domain (Fig. 2A). We defined *N*_res_ as the number of amino acid residues between the two labeled sites and *R*_E_ as the end-to-end distance between the fluorophores at these sites. We measured eighteen chain segments in COS-7 cells using our developed pipeline (see supplementary text for details). Briefly, our pipeline involved auto-segmentation to select the nuclear rim from each cell as a region of interest and extraction of measured donor fluorescence intensity profiles before and after acceptor photobleaching, defined as *I(t)* and *I’(t)*. By subtracting *I(t)* from *I’(t)*, the signals from the donor-only population and the background were eliminated, and the difference was taken as the pure FRET population. We directly observed the differences in FRET efficiencies among all mutants in the fluorescence intensity profiles (Fig. 2F). We detected a clear trend, where a smaller *N*_res_ showed a larger difference before and after acceptor photobleaching, indicating a higher FRET efficiency and smaller *R*_E_, and *vice versa*. To further compare the fluorescent lifetimes across cells, we converted individual fluorescence decay curves into a phasor plot — a graphical view enabling the cell-by-cell analysis of the complicated lifetime curves (Fig. 2G and Fig. S7B) (*40*). The phasor plot revealed not only some heterogeneity across cells but also an overall trend, where bigger *N*_res_ exhibited more left-shifted phasor value, corresponding to longer fluorescence lifetime and bigger *R*_E_. These results suggested that FLIM-FRET could be successfully used to spatially distinguish and map the chain dimensions of NUP98 FG-domain.

### Chain dimensions extracted from the polymer scaling law

We determined the chain dimensions of the NUP98 FG-domain *in situ* by means of the polymer scaling law that relates the root-mean-square end-to-end distance 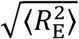 to the chain length *N*_res_ as given by (*34*)

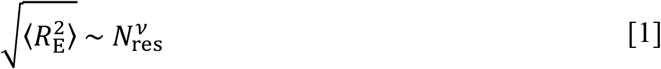

where *v* is the scaling exponent. In Flory’s homopolymer theory (*34*), *v* ∼ 0.3 indicates that the polymer is very compact, as self-interactions dominate over interactions with the “poor-solvent”; at *v* = 0.5 those interactions are balanced, while at *v* ∼ 0.6 the polymer is in “good-solvent” conditions and appears expanded. Flory’s theory can be further extended to describe the densely-grafted polymer brushes where *v* ∼ 1 (*41*). Note, that the scaling law is derived for infinitely long homopolymers. Despite its limiting definition (*42*), the law has been applied to calculate an apparent scaling exponent for finite-length protein. (*28, 43*). In a nutshell, the apparent scaling exponent captures a complex distance distribution in one number and thus provides excellent economy to describe changes in how protein conformational changes are tuned by their environment.

Here, we measured lifetime decay curves for the eighteen chain segments of NUP98 (Fig. S7A) and fitted them with the Gaussian chain model to extract the scaling exponent *v* (see supplementary text for details). We acknowledge that more sophisticated models exist, at the cost of having to use more fitting parameters. We obtained a scaling exponent *v* = 0.56 ± 0.05 (> 0.5), indicating that the probed NUP98 FG-domain adopted a rather extended conformation inside the NPC in the cells (Fig. 3). This result contrasts our *in vitro* single-molecule FRET (smFRET) measurements of the purified NUP98 FG-domain at a pico-molar concentration (Fig. S8), where the scaling exponent was *v* = 0.29 ± 0.05 (< 0.5) in line with other solution measurements (*28, 44*– *46*). With water acting as a ‘poor solvent’ for the hydrophobic side-chains, the single NUP98 FG-chain tends to become buried in a globule-like protein conformation *in vitro*. By contrast, the central channel of the NPC is enriched with other FG-NUPs, which collectively could provide a “good-solvent” environment to expand the polymer chains, reminiscent of the chain conformations in polymer melts (*34*). Note, that in the NPC also high quantities of other proteins such as nuclear transport receptors exist, which can contribute to the “good-solvent” conditions.

**Fig. 3.**
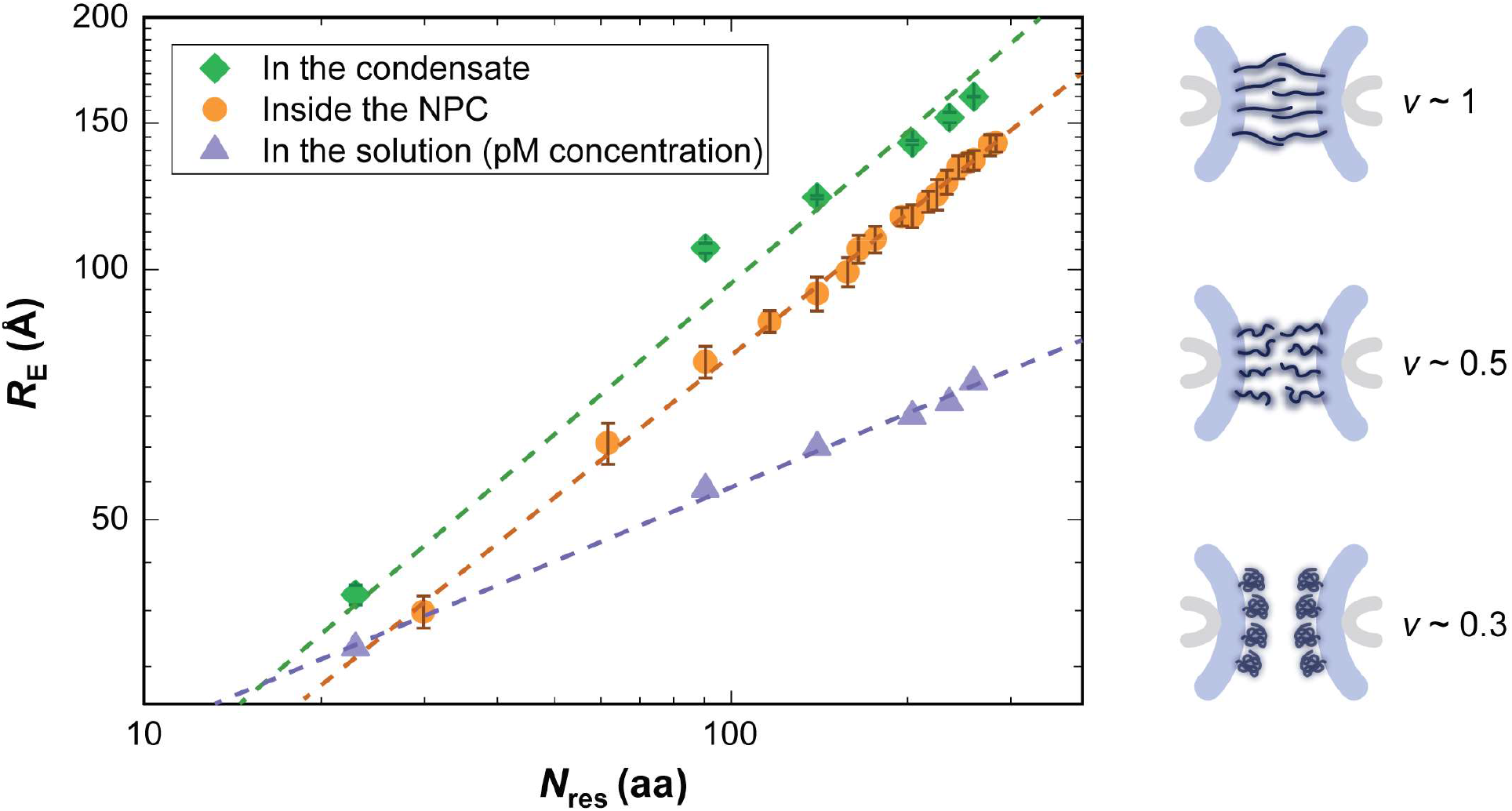
Scaling law of the NUP98 FG-domain. Left: NUP98 FG-domain showed more extended conformations inside the NPC (scaling exponent *v* = 0.56 ± 0.05) and in the phase-separated condensate (*v* = 0.60 ± 0.03). NUP98 FG-domain showed a more collapsed conformation in the physiological solution on a single-molecule level (*v* = 0.29 ± 0.03). Right: Schematics showing the conformations of FG-NUPs inside the NPC associated for different scaling exponents. When *v* ∼ 1, FG-NUPs behave like densely grafted polymer brushes. When *v* ∼ 0.5, FG-NUPs adopt an ideal-chain conformation. When *v* ∼ 0.3, FG-NUPs adopt a collapsed conformation.

### FG-NUP motions in NPC central channel revealed by coarse-grained modeling

The FLIM-FRET experiments empower MD simulations to provide us with a 3D view of the organization of FG-NUPs in functional NPCs. The disordered FG-domains are grafted on the NPC scaffold *via* short-folded domains. The recent structure of the human NPC scaffold (*16*) allows us to anchor FG-NUPs at the correct positions, orientations, and grafting densities. This enabled us to build a coarse-grained bead-spring polymer model (*47*) and perform MD simulations of the so-far elusive FG-domains attached to the NPC scaffold (see supplementary text). We first parameterized the effective NUP-NUP interaction strength as 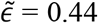 defined in supplementary text equation 17 by matching the phase behavior of *in vitro* reconstituted NUP98 FG-condensates that mimic the permeability barrier (*48*) (Fig. S13A, Movie S1). We also performed FLIM-FRET measurements on the freshly prepared NUP98 FG-droplets, which showed similar permeability barrier properties as functional NPCs (Fig. 1C). The measured distance distribution agreed well with the simulations (Fig. S13D), validating the parameterization approach. Also here, the measured scaling exponent of FG-condensates was *v* = 0.60 ± 0.03, in line with “good-solvent” conditions (Fig. 3, Fig. S9) and with the ability to function as permeability barriers. However, when the same interaction strength of 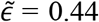 was applied to the whole-NPC simulations, the inner-ring FG-NUPs collapsed onto the scaffold to form surface condensates (Movie S2-S4), despite only weak direct interactions between FG-NUPs and the scaffold 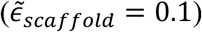. The surface condensates left a void at the center of the pore with a diameter of ∼20 nm, which does not seem in good agreement with the permeability barrier function of the NPC to block the unaided passage of large cargo. After a slight adjustment of the interaction strength to 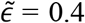, the FG-NUPs recovered the distance distribution matching our FLIM-FRET data (Fig. 4A). We observed that under such a condition, the FG-NUPs formed extended coil configurations and the inner-ring FG-NUPs fluctuated extensively to form a dynamic barrier across the central channel (Fig. 4B, Movie S3). Remarkably, the optimal FG-NUP interaction strength found in this way nearly coincides with the critical strength 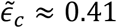 where condensation sets in. The large FG-NUP motions seen in the MD simulations thus amount to critical fluctuations that create a highly dynamic polymer network. These extensive and fast motions of individual molecular chains could provide an explanation for the rapid but selective molecular transport in the permeability barrier (*46, 49, 50*).

**Fig. 4.**
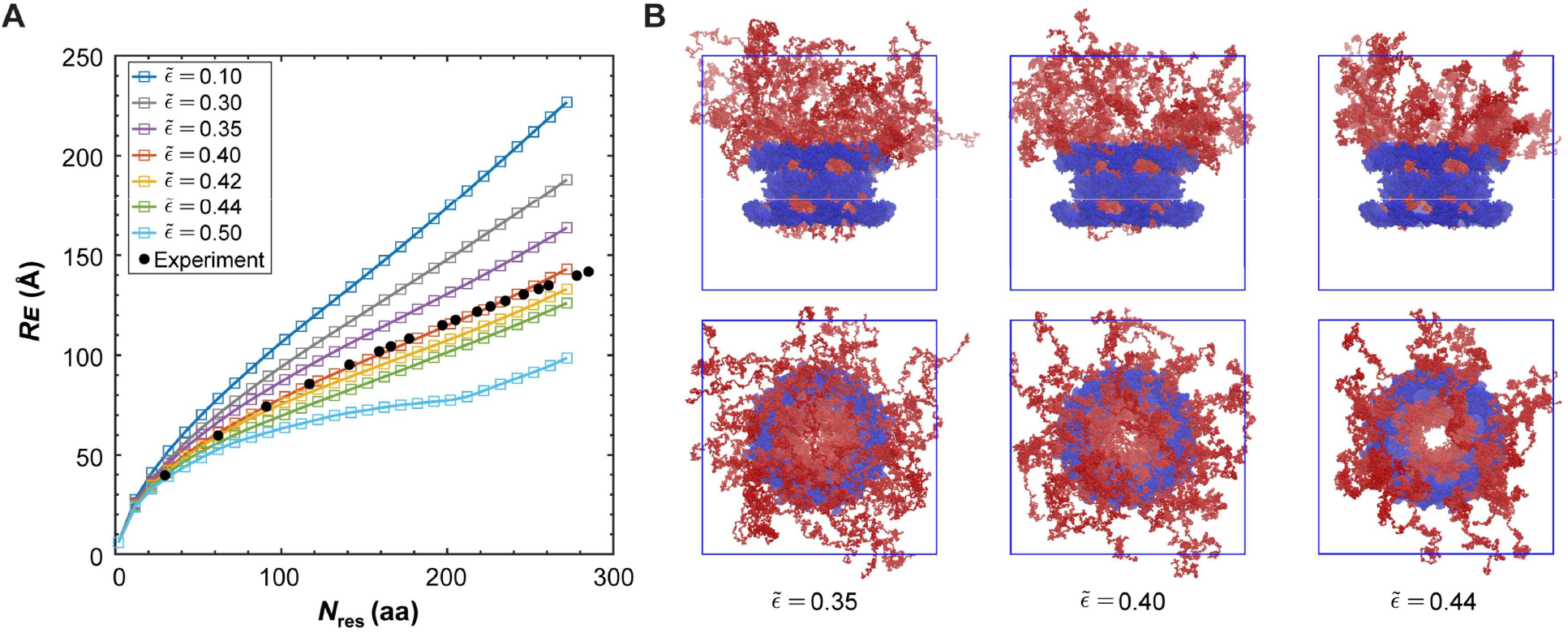
Coarse-grained MD simulations of FG-NUPs in the NPC. (**A**) Mean distances *R*_E_ of beads on the same NUP98 FG-domain in NPC as function of residue separation *N*_res_ with different effective NUP-NUP interaction strength 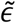. The distances from the FLIM-FRET experiments are shown as filled spheres, which overlaps with the simulation condition 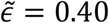. (**B**) Side and top views of the NPC at the end of the MD simulations as 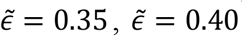, and 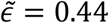 (scaffold: blue; FG-NUPs: red, see Movie S2-S4).

## Discussion

Here we developed an experimental approach using site-specific fluorescent labeling of IDPs in non-fixed mammalian cells and fluorescent lifetime imaging microscopy (FLIM) to directly decipher their plasticity *via* FRET in the functional state. We showed that this approach works for the sub-resolution permeability barrier of NPC, a nanocavity with a diameter of ∼ 60 nm, filled with ∼50 MDa of highly concentrated FG-NUPs. By measuring the end-to-end distances of different segments of the labeled FG-NUPs using FLIM-FRET, we obtained the distribution from which we could estimate an apparent scaling exponent and revealed that the intact NPC environment provides a “good solvent” where the FG-domains adopt extended conformations compared to the solution state *in vitro*.

We synergistically combined our FLIM-FRET measurements with computational modeling and investigated the conformational behavior and interaction mechanisms of FG-NUPs at the molecular level. We found in the MD simulations that a simple polymer model could capture the molecular dynamics of FG-NUPs and reproduce the FLIM-FRET right at the critical interaction strength, which defines the energetic threshold to protein condensate formation and is associated with large fluctuations in the dynamic polymer network (Movie S3). For weaker interactions, the polymer network was too loose to generate a polymer network (Movie S2). For stronger interactions, the FG-NUPs formed a surface condensate whose collapse onto the scaffold was driven by the geometry of the grafting points, not by the weak direct interactions with the scaffold (Movie S4). Indeed, the chain extension is exquisitely sensitive to the NUP-NUP interaction strength 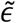 in the NPC (Fig. 4A), whereas in bulk condensates the extension is nearly insensitive (Fig. S13B). Thanks to the recent advances in NPC scaffold structure determination, the actual grafting points for the FG domains are now known with very high confidence (*13*–*20*). We conclude, first, that near-critical conditions, 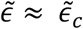, best capture the FLIM-FRET measurements; and second, that the bulk condensate, despite having similar permeability barrier properties as the intact NPC, is an incomplete approximation of the actual permeability barrier, whose materials properties are modulated by the anchoring of a distinct number of FG-NUPs with 3D precision on a half-toroidal NPC scaffold. In terms of nuclear transport selectivity, the consequences are significant: the surface condensate leaves a significant hole at the center; this hole is filled by the critical polymer mixture. Importantly, it is in the regime around the critical point that the FRET distance measurements are most discriminatory about the conformational state of NUP98 FG-domain (compact *versus* expanded). Remarkably, we also showed that the parameterization based on *in vitro* reconstitution studies failed to reproduce a functional pore, emphasizing the importance of applying the *in-situ* measurements in guiding the MD simulations. Our study removes much speculation about the conformational state of the FG-NUPs in the NPC and provides a sound coarse-grained model with amino acid precision to explain the function of the permeability barrier.

Computational modeling tools, like the most advanced AI-based AlphaFold, can very precisely predict structures of folded proteins, however, when applied to IDPs, they still largely require experimental constraints and validation (*51*). It is particularly necessary to acquire the constraints and validation from *in situ* experimental measurements, as we showed here, where a protein chain behaving globular-like *in vitro* can adopt a much more expanded conformation in the functional state due to the cellular environment. Our work serves as an exemplar of how the structural knowledge from cryo-electron tomography — which revealed the exact anchoring sites of IDPs — paired with *in situ* FLIM-FRET — which revealed the conformational state of the proteins inside the nano-sized NPC — can yield a complete coarse-grained picture of a cellular machinery enriched in IDPs inside the cell. The tools developed here are generally applicable to study the plasticity and functions of many other IDPs in the cell, filling a major technology gap in the field.

## Supporting information

Movie_S1_NUP98_FG-condensate_Eps_044

Movie_S2_NPC_Eps_035

Movie_S3_NPC_Eps_04

Movie_S4_NPC_Eps_044

Movie_S5_Single_NUP98_FG_chain_Eps_06

## Acknowledgments

We thank all the members of the Lemke laboratory for helpful discussions. We thank the Core Facilities of the Biology Faculty at Johannes Gutenberg University Mainz and the Protein Production Core Facility at Institute of Molecular Biology Mainz for expert assistance.

## Author contributions

E.A.L. and G.H. conceived the project. M.Y., S.M., P.S.T., S.M., H.R., and C.D.R. performed the experiments. M.H. and M.S. performed the simulations. M.Y. and M.H. analyzed the data and co-wrote the original draft together with G.H. and E.A.L. All the authors contributed to manuscript editing.

## Data and materials availability

All data are available in the main text or the supplementary materials. All plasmids can be obtained upon reasonable request.

