## Supplementary_Materials for "Deciphering the conformations and dynamics of FG-nucleoporins *in situ*"

M. Yu *et al.*

mailto:

**This PDF file includes:**

Materials and Methods
Supplementary Text
Figs. S1 to S13
Table S1 to S2
References (*52-75*)

**Other Supplementary Materials for this manuscript include the following:**

Movies S1 to S5

### Materials and Methods

Cell culture, transfection, and labeling

*Cell culture.* COS-7 cells were maintained in Dulbecco's Modified Eagle Medium (Life Technologies) supplemented with 10% v/v fetal bovine serum, 1% Pen-Strep, 1% L-glutamine (Sigma Aldrich), and 1% sodium pyruvate (Life Technologies) at 37 °C, 5% $CO_2$, and passaged every 2-3 days up to 15-20 passages.

*Cell transfection.* Cells were trypsinized and seeded into 35-mm imaging dish (Cat. No. 81158, ibidi) 24 hrs before transfection. The cells were transfected at a confluency of 60-70% with plasmids of interest listed in Table S1 (for Amber suppression with synthetic organelles, e.g., pcDNA3.1-hsNUP98$^{TAG}$-BoxB and pcDNA3.1-TOM20-FUS-$\lambda_{N22}$-PylRS(AF)-tRNA at a mass ratio of 1:1) using jetPRIME® according to manufacturer's protocol. After 4-5 hrs the medium was changed, and 10 mM HEPES with 50 µM {[(E)-cyclooct-2-en-1-yl]oxy}carbonyl)-L-lysine (TCO*A, SciChem, for in-cell labeling) or t-butyloxycarbonyl-lysine (BOC, Iris Biotech, for control measurements) were added, respectively.

*Cell labeling.* At 20-24 hrs post-transfection, COS-7 cells were washed twice with transport buffer (TB: 20 mM HEPES, 110 mM KOAc, 5 mM NaOAc, 2 mM $Mg(OAc)_2$, 1 mM EGTA, 2 mM DTT, pH 7.3) supplemented with PEG6000 (5 mg/mL) to avoid osmotic shock, and permeabilized for 2 min with 20 µg/mL digitonin (AppliChem) in TB. After the permeabilization, cells were washed twice with TB to remove digitonin and labeled with a dye solution containing 33.3 nM AZDye594-tetrazine (Click Chemistry Tools) and 66.6 nM LD655-tetrazine (Lumidyne technologies) in TB for 2 min. To remove residual dyes, the cells were washed in TB for 10 min at 37 °C three times. Cells were imaged immediately at room temperature. The lid was kept closed for the duration of cell imaging for 2 hrs.

Passive exclusion assay and active transport assay for labeled cells

*Passive exclusion assay.* COS-7 cells were transfected, permeabilized and labeled with 100 nM LD655-tetrazine as described above. After the last wash, the imaging dish was mounted on the custom-built confocal microscope. After gently removing the washing buffer, 0.5 µM 70 kDa FITC-Dextran (Sigma 53471) in TB was added to the dish, and the cells were imaged.

*Active transport assay.* To allow the import complex to form, 0.5 µM IBB-MBP-GFP cargo was first preincubated with 1 µM Importin-β on ice for 10 min, and then combined with the rest of the transport mix (5 µM RanGDP, 4 µM NTF2, and 2 mM GTP in TB). After that, the complete import mixture was added to the labeled and permeabilzied COS-7 cells as described above.

To ensure the NPC was functional during the 2-hr time window of FLIM measurements for each dish, passive exclusion assay and active transport reactions were performed for both the freshly labeled cells and the cells incubated in TB at RT for 2 hrs after the labeling steps.

Protein expression, purification, and labeling

*NUP98 FG-domain purification*. Homo sapiens NUP98 FG domain (1-505aa with Gle2-binding domain (GLEBS), a structured domain in between the two FG-domains, removed) was cloned into a pQE-14his-TEV vector. *E. coli* BL21 AI cells were grown in terrific broth (TB) medium containing 50 µg/ml of kanamycin at 37 °C, and protein expression was induced with 0.02% arabinose and 1 mM IPTG at $OD_{600}$ = 0.6. After 16 hrs of expression at 18 °C, cells were harvested by centrifugation and lysed in cell disruptor (CF1, Constant Systems) in the lysis buffer containing 6 M GdmCl, 0.2 mM Tris(2-carboxyethyl)phosphine (TCEP), 20 mM imidazole, and 1x PBS, pH 8. The lysate was centrifuged for 1 hr at 12000 g at 4 °C to remove cell debris, and the supernatant was incubated with Ni-beads for 2 hrs at 4°C. The Ni-beads with lysate were loaded in polypropylene tubes (Qiagen), washed twice with 2 M GdmCl and 20 mM imidazole, pH 8, and eluted with buffer containing 2 M GdmCl and 500 mM imidazole, pH 8. To remove the His-tag the elution was dialyzed in 0.5 M GdmCl and 50 mM Tris-HCl, pH 8, and cleaved overnight with TEV protease at room temperature. Proteins were incubated again with Ni-beads in 2 M GdmCl and 50 mM Tris-HCl, pH 8 to remove cleaved His-tags, TEV protease, and non-specific proteins with Ni-bead affinity. The flow-through containing the NUP98 FG domain was collected and further purified by size exclusion chromatography (Superdex 200, Akta pure protein purification system, Cytiva) in 2M GdmCl, 0.2 mM TCEP, and 1x PBS, pH 8. Fractions were analyzed by SDS-PAGE and stained with Coomassie blue. Pure fractions were pooled and concentrated to around 15 mg/mL in 4M GdmCl using 3 kDa MWCO centrifugal filters (Amicon, Merck), with the concentration measured by BCA protein assay kit (Thermo Fisher). The proteins were flash-frozen and stored at -80 °C.

*NUP98 FG-domain labeling in vitro*. For labeling, the purified NUP98 FG-domain with single or double cysteine mutations was exchanged to 4 M GdmCl, 1x PBS, 0.1 mM EDTA, 0.2 mM TCEP, pH 7. Labeling with Alexa Fluor 594 maleimide (Thermo) and LD 655 maleimide

(Lumidyne technologies) was done at the molar ratio of 1:2 (dye : protein) overnight at 4 °C. The reaction was quenched with 10 mM DTT in 4 M GdmCl, 1x PBS, pH 7. Unreacted dye was washed off using 3 kDa MWCO centrifugal filter (Merck Millipore), and the labeled protein was further purified with Superdex 200. Pure fractions were chosen, pooled, and concentrated as described above, and the final concentration was measured by absorbance spectrometer Duetta (Horiba). The proteins were flash-frozen and stored at -80 °C.

FLIM-FRET imaging setup

The custom-built FLIM-FRET imaging setup was equipped with picosecond pulsed laser diode heads including the wavelengths of 485 nm (LDH-D-C-485, PicoQuant), 560 nm (LDH-D-TA-560, PicoQuant), and 660 nm (LDH-D-C-660, PicoQuant). The laser heads were controlled through a multichannel picosecond diode laser driver (Sepia II PDL 828, PicoQuant). The beams were coupled into a single-mode polarization-maintaining optical fiber (KineFLEX-P-2-S-405/640-2.5-2.5-p2) and fiber coupler (60FC-4-RGBV11-47. Schäfter + Kirchhoff). The beam traveled through a Glan-laser polarizer (Thorlabs) and was directed into a laser scanning system (FLIMbee, PicoQuant). The three galvo mirrors in the scanning system were imaged onto the backfocal plane of the objective (60x SR Plan Apo IR, 1.27 NA, Nikon) with 200 mm tube lens. The fluorescence emission was focused onto a pinhole (100 μm for cell measurements, 50 μm for droplet measurements, and 100 μm for single-molecule FRET measurements), and then separated into parallel and perpendicular components using a 50/50 polarizing beam splitter (Thorlabs). Each component was further separated by two sets of beamsplitters (ZT561 RDC and T647 LPXR, Chroma), passed through three sets of bandpass filters (green channels: 525/50 BrightLine HC; orange channels: 609/57 BrightLine HC, Semrock; red channels: ET700/75m, Chroma), and focused onto the single-photon counting detectors (green channels and orange channels: PMA Hybrid 40, PicoQuant; red channels: τ-SPAD, PicoQuant). The signals from the photon detectors were recorded by a TCSPC system (HydraHarp 400, PicoQuant) at a time resolution of 16 ps. Data acquisition was carried out with SymPhoTime 64 software (PicoQuant).

Determination of Förster distance $R_0$

The Förster distance $R_0$ is the distance between a pair of fluorophores at which the FRET efficiency is 50%, which was calculated as,

$$R_0 = 0.211 \sqrt[6]{\frac{\kappa^2 \phi_D J(\lambda)}{n^4}} \ (R_0 \text{ in Å}) \quad [1]$$

where $\kappa^2$is the orientation factor, $n$ is the refractive index ($n$ = 1.375 for cell measurements (*52*), and $n$ = 1.52 when imaging droplets sitting on the coverslip (from manufacturer's datasheet)), $\phi_D$ is the quantum yield of the donor without energy transfer, and $J(\lambda)$ is the overlap integral between the donor emission and acceptor absorption spectra at wavelength $\lambda$, given by,

$$J = \int F_D(\lambda)\varepsilon_A(\lambda)\lambda^4 \mathrm{d}\lambda \,/ \int F_D(\lambda)\mathrm{d}\lambda \quad [2]$$

where $F_D(\lambda)$ is the radiation emission intensity of the donor at wavelength $\lambda$, and $\varepsilon_A(\lambda)$ is the extinction coefficient of the acceptor. The emission spectra of the donor and the excitation spectra scan of the acceptor were measured on a Leica SP8 STELLARIS microscope using xyλ or xyΛ acquisition mode, for single-labeled cells with the donor or the acceptor, respectively. The acquired spectra were then compared with the spectra measured for free dyes in TB using the absorbance spectrometer (Duetta) and no spectral shift was detected. $\kappa^2$ can be assumed to be ~2/3 when the dyes can freely rotate, which is supposed to be the case as all our dyes have a C5 flexible linker between the conjugating group and the chromophore. We also measured the anisotropies for the sample labeled with donor or acceptor dye inside the NPC, in the FG-condensates *in vitro*, and in the physiological buffer on a single-molecule level. The fundamental anisotropies were found to be smaller than 0.3 in all cases, and the error in the distance was proved to be below 10% (*39*). The $R_0$ for in cell measurements was determined as 77.1 Å, and for *in vitro* condensates measurements as 74.1 Å.

FLIM-FRET for cell measurements

The average power of laser excitation was optimized to collect enough photons from the cell within a reasonable time but avoid photon pile-up and other artifacts in the fluorescence lifetime measurements. The instrument response function (IRF) was measured on a daily basis using a freshly prepared saturated solution of KI and Erythrosine B (*53*). The temporal offsets of the parallel and perpendicular detectors were pre-aligned by the measured IRFs. The cell measurements were performed using the following imaging settings: the pixel size of 100 nm, the image size of 256 × 256 pixels, the pixel dwell time of 150 μs, and the time resolution of 16 ps.

The fluorescence photons were detected in T3 mode and collected from the perpendicular and parallel detectors for each color individually.

To measure FLIM-FRET, the morphology of the labeled cell was first checked with 660 nm laser excitation to ensure that no GLFG bodies (normally a sign of overexpressed NUP98 in the cell (*54*)) existed in the nucleus. The acceptor intensity per pixel of the nuclear rim was then checked with 660 nm laser excitation to further assess the expression level of the mutant NUP98, ensuring that the endogenous population of NUP98 remained substantially higher than the exogenous one. Specifically, we determined such an acceptor intensity range based on the criteria that, on the nuclear rim, the relative FRET efficiency (i.e., the proximity ratio, $E_{\text{rel.}} = \frac{I_{\text{A}}}{I_{\text{A}}+I_{\text{D}}}$, $I_{\text{A}}$ and $I_{\text{D}}$ are the total acceptor and donor fluorescence intensities, respectively, excited by 560 nm laser) did not corelate with the acceptor intensity per pixel excited by 660 nm laser (Fig. S6). This intensity threshold was further verified by acceptor photobleaching assays on cells that expressed single-Amber-mutant NUP98 and labeled with donor and acceptor dye mixture. The average fluorescence lifetime of the donor dye did not change before and after acceptor photobleaching as shown in (Fig. 2B and D), indicating no inter-molecular FRET could be detected.

After checking the expression level of the mutant NUP98 with 660 nm laser excitation, we imaged the selected cell for 5 min using 560 nm laser excitation with an average power of 40 μW at 40 MHz, and then for 30 sec using 660 nm laser excitation with an average power of 35 μW at 40 MHz. Next, the acceptor labeling was photobleached using 660 nm laser excitation with an average power of 300 μW at 40 MHz for 2 min. The donor signal was measured again post-photobleaching for 5 min using 560 nm laser excitation with an average power of 40 μW at 40 MHz.

The recorded images were processed using an automatic segmentation pipeline developed based on the software package PAM in Matlab (*55*). The nuclear rim was selected as a region of interest (ROI) using a thresholding algorithm according to the intensity and average lifetime of each pixel. The time-resolved donor fluorescent intensity profiles before and after acceptor photobleaching were extracted from the selected ROI, respectively. The total fluorescence decay was calculated by combining the parallel and perpendicular fluorescence decays ($I_{\parallel}$ and $I_{\perp}$),

$$I(t) = (1 - 3L_2)GI_{\parallel}(t) + (2 - 3L_1)I_{\perp}(t) \quad [3]$$

where $L_1$ and $L_2$ are factors accounting for polarization mixing caused by the high numerical aperture objective lens, and $G$ is the factor accounting for the difference in the detection efficiencies $\eta$ between parallel and perpendicular polarization, given by,

$$G = \frac{\eta_{\perp}}{\eta_{\parallel}} \quad [4]$$

$G$ factor was determined as a ratio of the average intensities of the perpendicular and parallel donor channels when measuring a solution of Tris(2,2'-bipyridyl)dichlororuthenium(II) chloride ($[Ru(bpy)_3]Cl_2$). For each mutant, the total fluorescence decays for ~100 cells were added up for further fitting analysis (see SI text for details).

*In vitro* droplet assay and FLIM-FRET measurements

The purified and labeled NUP98 FG-domain was mixed with unlabeled protein at a molar ratio of 1:5000 in 2 M GdmCl, 1x PBS, pH 7. 1 µl of such a mixture was then quickly mixed with 24 µl TB supplemented with 5 mg/mL PEG6000 in a chambered coverslip (Cat. No. 81507, ibidi) and imaged immediately with the custom-built confocal microscope. The final total concentration of NUP98 in the system was 10 µM for unlabeled and 2 nM for labeled. The trace amount of GdmCl left in solution was below < 100mM (a negligible level). Note, that here we removed the structured GLEBS domain because it could potentially misfold and trigger aggregation when rapidly changing the buffer condition from denaturing to native in the above-described droplet assay.

For the passive exclusion assay and the facilitated/active transport assay on the phase-separated condensates, unlabeled NUP98 and LD655-labeled one were mixed with the ratio described above, to avoid the crosstalk with the fluorescently labeled cargos (i.e., FITC or GFP). After mixing the protein with TB for 5 min, the buffer was carefully replaced by 0.5 µM 70 kDa FITC-Dextran in TB or 0.5 µM IBB-MBP-GFP in the transport mixture as described for the labeled cells avoiding disturbing the condensates on the coverslip surfaces and imaged immediately.

To measure FLIM-FRET on the NUP98 FG-condensates, we focused on the first 5 min as the condensates behaved more liquid-like during this time (*25*) (e.g., FG-droplets merged quickly as shown in Fig. S2) . We applied the same conformation distribution model (i.e., the Gaussian chain model) to fit the lifetime curves of the FG-condensates as we used for in cell measurements.

The formed condensates were measured for 5 min using 560 nm laser excitation with an average power of 70 μW at 40 MHz, and the imaging settings were as follows: the pixel size of 200 nm, the image size of 256 × 256 pixels, the pixel dwell time of 100 μs, and the time resolution of 16 ps. The fluorescence photons were detected in T3 mode and collected from the perpendicular and parallel detectors for each color individually. The FG-droplets were selected as the ROI based on the intensity and average lifetime in SymPhoTime 64 software. The time-resolved donor fluorescent intensity profiles were extracted from the ROI pixels for analysis.

To ensure no inter-molecular FRET was detected, single-cysteine-mutated NUP98 was labeled with either the donor dye or with the mixture of donor and acceptor dyes. The labeled protein was then mixed with the unlabeled protein with the same concentration as described above to perform the droplet assay. The donor fluorescence lifetimes of the donor-only and the donor-acceptor mixture showed no difference, indicating that no inter-molecular FRET could be detected (Fig. S9).

Phasor transform of FLIM data

To analyze the lifetime decays on a single-cell basis, the raw intensity profile $I(t)$ of each selected nuclear rim was plotted as a single point in the phasor plot by applying the Fourier transform to the measured decay data, given by

$$g(\omega) = \frac{\int_0^T I(t)\cos(2\pi f t)\mathrm{dt}}{\int_0^T I(t)\,\mathrm{dt}} \quad [5]$$

$$s(\omega) = \frac{\int_0^T I(t)\sin(2\pi f t)\mathrm{dt}}{\int_0^T I(t)\,\mathrm{dt}} \quad [6]$$

Where $g(\omega)$ and $s(\omega)$ are the $x$ and $y$ coordinates of the phasor plot, respectively, $f$ is the repetition rate of laser excitation, i.e., 40 MHz, and $T$ is the repeat frequency of the acquisition. To establish the correct scale for the plotted phasor points, the coordinates of the phasor plot were first calibrated by applying Fourier transform to the measured IRF trace and setting it as the zero lifetime (*56*). Each phasor point from the acquired FLIM data was then calibrated accordingly using the same calibration parameters so that the final phase plot was referenced relative to the calibration standard. The above procedure was performed with self-written code in MATLAB.

Single-molecule FRET measurements

Single-molecule FRET measurements were performed with the custom-built confocal microscope mentioned above. The sample was illuminated with 560 nm and 660 nm lasers in pulsed interleaved excitation (PIE) mode at a repetition rate of 32 MHz. Photon counts were recorded with a resolution of 16 ps. Purified and double-labeled NUP98 was diluted to the final protein concentration of 50 pM in 1x PBS supplemented with 10 mM fresh DTT. The donor dye was excited with 560 nm laser at an average power of 70 μW and 660 nm laser at an average power of 20 μW. Data acquisition was carried out with SymPhoTime 64 software.

Acquired data was analyzed with PAM software package (*55*) for burst search and the identified bursts were further analyzed in the BurstBrowser. The FRET efficiency $E$ in a burst is defined as,

$$E = \frac{I_A^D}{\gamma I_D^D + I_A^D} \qquad [7]$$

and the stoichiometry $S$ is defined as,

$$S = \frac{\gamma I_D^D + I_A^D}{\gamma I_D^D + I_A^D + I_A^A} \qquad [8]$$

where $I_A^D$ is the acceptor fluorescence detected upon donor excitation, $I_D^D$ is the donor fluorescence upon donor excitation, $I_A^A$ is the acceptor fluorescence detected upon acceptor excitation, and $\gamma$ is the factor accounting for the detection efficiency of the acceptor and donor channels. After performing the correction for the leakage of donor fluorescence into the acceptor channel ($\alpha = 1.00$) and direct acceptor excitation ($\delta = 0.30$) and confirming minimal variation of quantum yields among different mutants, the $\gamma$ parameter was extracted from $E_{app}$ and $S_{app}$ (*57*). A linear fit to a plot of $1/S_{app}$ versus $E_{app}$ yields intercept $a$ and slope $b$ which relates to $\gamma$ in the following way,

$$\gamma = (a - 1)/(a + b - 1) \qquad [9]$$

We determined the $\gamma$ parameter of labeled NUP98 in 1x PBS to be 2.51. Examples of single-molecule transfer efficiency histograms are shown in Fig. S8. The FRET population was selected and fitted with a 2D bi-gaussian function due to high FRET efficiency. The fitted values were taken as the average FRET efficiency, which is defined as,

$$\langle E\rangle = \int_0^\infty E(R)\rho(R)\,\mathrm{d}R \tag{10}$$

where $\rho(R)$ describes the distance distribution of the end-to-end distance. A typical smFRET experiment does not contain enough information to retrieve a model-free distance distribution, and one must therefore choose a model to fit. Here we adopt the Gaussian chain model, which assumes that the monomers occupy zero volume and excluded volume effects are not considered. Despite its simplicity, it is commonly used for the analysis of IDPs (*28*, *58*). The distance distribution function takes the form given by

$$\rho(R) = 4\pi R^2 \left(\frac{3}{2\pi\langle R_E^2\rangle}\right)^{\frac{3}{2}} e^{-\frac{3R^2}{2\langle R_E^2\rangle}} \tag{11}$$

where $\sqrt{\langle R_E^2\rangle}$ is the root mean squared end-to-end distance. By comparing the fitted $\langle E\rangle$ with the calculated average FRET efficiency derived from the model, $\sqrt{\langle R_E^2\rangle}$ was obtained for each mutant.

### Supplementary Text

FLIM analysis methods

The fluorescence lifetime decays for the eighteen chain segments of NUP98 inside the NPC were extracted from the selected ROIs using the FLIM-FRET measurements pipeline described above. The observed emission from the donor channels consists of three populations: donor-only population $I_{\mathrm{Donly}}$, FRET population between donor and acceptor dyes $I_{\mathrm{FRET}}$, and cellular background signal $I_{\mathrm{bg}}$. The time-resolved fluorescence intensity in the donor channel can be described by

$$\begin{aligned} I(t) &= \left[I_{\mathrm{Donly}}(t) + I_{\mathrm{FRET}}(t) + I_{\mathrm{bg}}(t)\right] \otimes IRF \\ &= \left\{A_{\mathrm{D}} e^{-\frac{t}{\tau_D}} + A_{\mathrm{FRET}} \int_0^\infty \rho(R) e^{-\frac{t}{\tau_D} * [1+\left(\frac{R_0}{R}\right)^6]} \mathrm{d}R + A_{\mathrm{bg}} \sum_{i=1}^{N} \alpha_i\, e^{-\frac{t}{\tau_{bg_i}}}\right\} \otimes IRF \end{aligned} \tag{12}$$

$$A_{\mathrm{D}} + A_{\mathrm{FRET}} + A_{\mathrm{bg}} = 1 \tag{13}$$

where $A_{\mathrm{D}}$, $A_{\mathrm{FRET}}$, and $A_{\mathrm{bg}}$ are the species fraction, i.e., the initial intensities (at $t = 0$) for the three components, respectively, $\tau_D$ is the donor fluorescence lifetime in the absence of an energy transfer acceptor, $\rho(R)$ is the distribution of the end-to-end distance that characterizes the conformational plasticity of the chain segment, $R_0$ is the Förster distance of the dye pair, $\tau_{bg_i}$ is the lifetime characterizing the particular exponential components of the cellular background with the species fraction $\alpha_i$, and IRF is the instrument response function. The ensembled lifetime of the cellular background was determined in ~ 200 cells with Amber-mutant-NUP98 expressed in the presence of Lys-BOC, a nonreactive ncAA, and the added-up lifetime decay was fitted with a three-exponential function with TauFit in PAM. After acceptor photobleaching, the time-resolved fluorescence intensity in the donor channel can be described by

$$I'(t) = \left\{ A_{\mathrm{D}} e^{-\frac{t}{\tau_D}} + A_{\mathrm{FRET}} e^{-\frac{t}{\tau_D}} + A_{\mathrm{bg}} \sum_{i=1}^{N} \alpha_i \, e^{-\frac{t}{\tau_{bg_i}}} \right\} \otimes IRF \qquad [14]$$

Assuming no photobleaching of the donor and background during the FLIM-FRET measurements before and after acceptor photobleaching, the difference in the intensity is calculated by subtracting Eq. 12 from Eq. 14,

$$I'(t) - I(t) = A_{\mathrm{FRET}} \left\{ e^{-\frac{t}{\tau_D}} - \int_0^{\infty} \rho(R) e^{-\frac{t}{\tau_D} * [1 + \left(\frac{R_0}{R}\right)^6]} \mathrm{d}R \right\} \otimes IRF \qquad [15]$$

where the signals from the donor-only population and the background were eliminated, and the difference was only from the FRET population. We have noticed that the signals from the donor and the background would bleach during the FLIM-FRET measurements. Therefore, we obtained the bleaching rate $\sigma$ experimentally and corrected the intensity difference as $I'(t) - \sigma I(t)$. Next, we normalized the intensity profiles after the acceptor photobleaching and, using the same scaling factor for the corresponding intensity profiles before the acceptor photobleaching, to qualitatively demonstrate a trend in Fig. 2F, in which the higher FRET sample showed a bigger difference in the FRET efficiency across all mutants.

The stoichiometric labeling method that we adopted here results in an inherent donor-only component, which decreases the resolution of the distance distribution due to its distinct decay kinetics and higher emission intensity (*39*). To reduce the donor-only component, we optimized the stoichiometric ratio between donor and acceptor dye in the mixture, and found that donor:acceptor = 1:2 offered us a relatively larger FRET population to increase the lifetime resolution as well as enough photon counts from the donor channel within a reasonable measuring time. It has been shown that it is practical to recover correct values of distance distribution parameters if the FRET labeling ratio $L = A_{\mathrm{FRET}}/(A_{\mathrm{FRET}} + A_{\mathrm{D}}) > 0.5$, which was satisfied with our optimized stoichiometric labeling ratio (*59*). The fitting accuracy is further improved if the ratio of the background is known, which we determined by performing acceptor photobleaching.

To extract the polymer scaling law for the eighteen chain segments of NUP98 inside the NPC, we performed a global fitting on the ensembled lifetime decays. To reduce the number of parameters to fit, we adopted the Gaussian chain model (Eq. 11), arguably the simplest and most cited model to describe the distribution of the end-to-end distance with least free fitting parameters, and embedded it into the self-written code adapted from the TauFit function of PAM. The lifetime decays after acceptor photobleaching were first fitted with a single-exponential function, where the background profile was fitted with the control measurements using a non-reactive ncAA BOC, as discussed above. The 95% confidence intervals for the fitted donor lifetime and the ratio of the background were obtained for each mutant from the Jacobian matrix. By substituting the polymer scaling law which is given by

$$\sqrt{\langle R_E^2 \rangle} = \rho_0 N_{\mathrm{res}}^{v} \quad [16]$$

and Eq. 11 into Eq. 12, the lifetime decay of each mutant could be directly expressed with the scaling exponent $v$, prefactor $\rho_0$, and the number of amino acid residues between the double-labeled sites $N_{\mathrm{res}}$. To perform a global fitting on the FLIM-FRET lifetime decays before acceptor photobleaching, $v$, $\rho_0$, and $A_{\mathrm{D}}$were set as global parameters, and the lower and upper boundaries of the donor lifetime and the background for each mutant were adopted from the 95% confidence intervals determined above. The scaling exponent was extracted from the global fitting using the Maximum Likelihood Estimator (MLE). To estimate the biological heterogeneity, the standard errors were calculated using the bootstrap resampling method.

To verify the fitting algorithm, we extracted the fluorescence intensity profiles of the detected bursts from the smFRET measurements dataset of the purified NUP98 in PBS *in vitro*. We then performed the lifetime fitting for each mutant with Eq. 11 and 12. The correlation of the $\sqrt{\langle R_E^2 \rangle}$ extracted from the lifetime fitting versus the $\sqrt{\langle R_E^2 \rangle}$, obtained from the smFRET analysis described above, is shown in Fig. S10 with $R^2 = 0.97$, validating the approach.

To extract $\sqrt{\langle R_E^2 \rangle}$ of different mutants from the lifetime decays of the NUP98 FG-condensates *in vitro*, a similar fitting pipeline as for in cell measurements was applied with some modifications. Since the phase-separated condensates demonstrated liquid-like behavior at the early stage (as we measured for the first 5 min) where the fluorescence does quickly recover after photobleaching, we could not perform the acceptor photobleaching to estimate the background signal ratio as we did for the cell measurements. Therefore, we first measured the lifetime profile for the condensates formed by the unlabeled NUP98 using the differential interference contrast (DIC) module to check the focusing plane. We then measured the lifetime decay for the unlabeled NUP98 mixed with the donor-only labeled sample with the same donor concentration as for the FRET samples and fitted the decay curve with a single-exponential function to determine the ratio of the background signal. After that, we performed the Gaussian chain model fitting of the lifetime curves for the FRET samples and extracted $\sqrt{\langle R_E^2 \rangle}$ for each mutant in the FG-condensates.

Molecular dynamics simulation methods

We performed molecular dynamics simulations to model the large-scale conformations and motions of FG-NUPs in the NPC and in the *in vitro* reconstituted NUP98 FG-condensates. To ensure efficient sampling, we used a coarse-grained polymer model, in which each amino acid is represented by a single bead without local structure.

*Composition and structure.* The model complements the recently resolved NPC scaffold of the human NPC (*16*) with a nearly complete set of FG-NUPs (see Table S2). The FG-NUPs were grafted onto the NPC scaffold in the constricted state (PDB ID: 7R5K) at established positions as indicated.

*Energy function.* We used a FENE potential ($U_{FENE}$) (*60*) to describe the disordered FG-NUPs, Lennard-Jones (LJ) interactions ($U_{LJ}$) between the CG beads, and a potential confining NUP98 chains. The total potential energy is

$$
\begin{aligned}
U &= U_{LJ} + U_{FENE} + U_C \\
&= 4k_BT \sum_{i<j, r_{ij}<r_c} \tilde{\epsilon}_{ij} \left[ \left(\frac{\sigma}{r_{ij}}\right)^{12} - \left(\frac{\sigma}{r_{ij}}\right)^{6} \right] + k_BT \sum_{\langle i,j=i+1\rangle, r_{ij}<2^{\frac{1}{6}}\sigma} \left[ \left(\frac{\sigma}{r_{ij}}\right)^{12} - \left(\frac{\sigma}{r_{ij}}\right)^{6} \right] \\
&\quad - \sum_{\langle i,j\rangle} 0.5 k_F R_0^2 \ln\left[1 - \left(\frac{r_{ij}}{R_0}\right)\right] + k_c \sum_{i \in Nup98, r_{xy,i}>R_c} \left(r_{xy,i} - R_c\right)^2
\end{aligned}
\qquad [17]
$$

The length and interaction energy scales of the LJ potential between a pair of beads $i$ and $j$ are $\sigma$ and $\tilde{\epsilon}_{ij}$, respectively. The latter is normalized by the thermal energy $k_BT$, where $k_B$ is the Boltzmann constant and $T$ is the system temperature. We fixed the interaction strength between scaffold and FG beads at $\tilde{\epsilon}_{ij} = \tilde{\epsilon}_{\text{scaffold}} = 0.1$, which ensures that chains do not stick to the scaffold. The interaction strength between FG-beads is given uniformly by $\tilde{\epsilon}_{ij} = \tilde{\epsilon}$. The sum over $\langle i, j = i + 1\rangle$ extends over the bonded beads of FG-NUPs. The bond contraction and stretching are controlled by the repulsive LJ and logarithmic terms, whose maximum bond limit is $R_0 = 1.5\sigma$ with $k_F = 80k_BT$. To mimic the membrane envelope underneath the scaffold, we applied an axial confinement on NUP98-FG beyond a radius of $R_c = 81\sigma \equiv 48.6$ nm, where $r_{xy} = \sqrt{x^2 + y^2}$ and $k_c = 80k_BT$.

*MD simulations*. We used the LAMMPS (*61*) to simulate the polymeric systems. The systems were thermalized with a Langevin thermostat (*62*) (at $k_BT = 1$) with a damping coefficient of $10\tau$. We used a uniform mass $m$ for all monomers and a characteristic time scale $\tau = \sqrt{ma^2/k_BT}$ . The time step of all simulations was set to $0.01\tau$ for the single-chain and condensate simulations and $0.001\tau$ for NPC simulations. We used block averaging with four non-overlapping blocks to estimate uncertainties. In the MD simulations of the FG-NUPs attached to the NPC scaffold, the residues of the FG-NUPs grafted to the scaffold (Table S2) and the scaffold itself were kept frozen.

We determined the bond length $\sigma$ by matching the end-to-end distance distribution of the CG model of single NUP98 FG-chain (1-499) to that in the Martini 2.2 level of coarse-graining (*63*, *64*). With a focus on the geometric extensibility, we worked in the limit of weak non-bonded interactions between distant amino acids. For the Martini simulations of the single chain, we used $\alpha$-scaling ($\alpha = 0.1$) (*65*) and for the CG model a weak cohesive strength between FG-beads ($\tilde{\epsilon} = 0.1$). The Martini MD simulations were performed with GROMACS 2020.6 (*66*, *67*). The first 499 N-terminal amino acids of NUP98 were converted into a Martini coarse-grained model using marinize.py python script (*63*, *64*). A cubic simulation box of size 30 nm was built and solvated with coarse-grained Martini water and 10 percent anti-freezing water. Ions were added to neutralize the system. The system was initially energy minimized and then equilibrated for 500 ns in NVT and NPT ensembles, respectively, using the velocity scaling thermostat (*68*) and the Berendsen barostat (*69*) at temperature $T = 300$ K and pressure $P = 1$ atm. The time constant of the thermostat was 1 ps, and that of the barostat 5 ps with a compressibility of $3 \times 10^{-4}$ bar$^{-1}$. For a 10-$\mu$s long production run in the NPT ensemble, we used a velocity scaling thermostat (*68*) and a Parrinello-Rahman barostat (*70*) with time constants of 1 ps and 12 ps, respectively. As shown in Fig. S11B, the probability distribution of the end-to-end distance $R_{ee}$ of the Martini model peaks at the same position as that of the CG model with $\sigma \equiv 0.6$ nm. In the following, we fix the FENE bond length at $\sigma \equiv 0.6$ nm.

In a second step, we adjusted the interaction energy between the FG-beads ($\tilde{\epsilon}$) by matching the model to the measured thermodynamic properties of NUP98 FG-chain (1-499) condensate formation (*71*, *72*). As shown in Fig. S13A, 500 chains were simulated inside a simulation box size of $360 \times 90 \times 90\ \sigma^3$. We adjusted the interaction strength $\tilde{\boldsymbol{\epsilon}}$ to match the experimentally measured concentration of the condensate (*71*, *72*) (see Fig. S13B). We note that in this way, we also get a good fit to the measured concentration of the dilute phase and thus the transfer free energy. The simulations of the condensate and NPC systems were performed for durations exceeding $1.8 \times 10^6\tau$ and $6 \times 10^4\tau$, respectively. For the analysis of condensate results, we used the last four blocks of $2 \times 10^4\tau$ each, and for the analysis of NPC results, we used the last four blocks of $1 \times 10^4\tau$ each.

*Comparison to smFRET and FLIM-FRET experiments.* We calculated the mean distances between the labeled sites across NUP98 FG-chains and simulation trajectories to compare to the smFRET and FLIM-FRET experiments. As in the experiments, we kept one site fixed (amino acid position 221 for NUP98 in the NPC) and swept across the C-terminal residues. For an isolated NUP98-FG (1-499) chain, the results are shown in Fig. S11C. We found that the experiments are best explained with a NUP-NUP interaction strength of $\tilde{\epsilon} = 0.6$, which is in the collapsed state (Movie S5). According to the radius of gyration of single chains (Fig. S11A), the midpoint of the coil-globule transition is at $\tilde{\epsilon} = 0.44$. We conclude that isolated NUP98 FG-chains in aqueous solution are collapsed.

*Flory-Huggins theory.* We used the mean-field Flory-Huggins (FH) model (*73–75*) to estimate the phase-coexistence curve and phase diagram from condensate simulations. The free energy of mixing per monomer is given by

$$\frac{\Delta \bar{F}_{mix}}{k_B T} = \frac{\phi}{N} \ln \phi + (1 - \phi) \ln(1 - \phi) + \chi\phi(1 - \phi) \quad [18]$$

where $\phi$ is the polymer volume fraction, $N = 499$ the number of monomers per chain, and $\chi$ the FH parameter quantifying the interaction energy per monomer. The first two terms account for the entropy of mixing chains and solvent, respectively, and the third term accounts for the enthalpy of mixing. For the interaction strength larger than the critical interaction strength, $\chi > \chi_c = \frac{1}{2} + N^{-1/2}$, the system separates into two coexisting dilute and dense phases with volume fractions of $\Phi'_{\tilde{\epsilon}}$ and $\Phi''_{\tilde{\epsilon}}$, respectively. The chemical potential of the system is determined by

$$\frac{\mu}{k_B T} = \frac{1}{k_B T}\frac{\partial \Delta \bar{F}_{mix}}{\partial \phi} = \frac{1}{N}\ln\phi - \ln(1-\phi) + \frac{1}{N} - 1 + \chi(1-2\phi) \qquad [19]$$

By setting the chemical potentials of the dilute and dense phases equal, and solving for $\chi$, one finds

$$\chi(\tilde{\epsilon}) = \frac{\frac{1}{\mathrm{N}}\log\left(\frac{\Phi''_{\tilde{\epsilon}}}{\Phi'_{\tilde{\epsilon}}}\right) + \log\left(\frac{1-\Phi'_{\tilde{\epsilon}}}{1-\Phi''_{\tilde{\epsilon}}}\right)}{2\left(\Phi'_{\tilde{\epsilon}} - \Phi''_{\tilde{\epsilon}}\right)} \approx A\tilde{\epsilon} + B \qquad [20]$$

We find that for the volume fractions $\Phi'_{\tilde{\epsilon}}$ and $\Phi''_{\tilde{\epsilon}}$ observed in the simulations, to a good approximation $\tilde{\epsilon}$ and $\chi$ are linearly related with coefficients A=1.773 and B=□0.1938 (Fig. S12). To interpret our simulation data in the FH model, we thus determined the volume fraction of each phase using $\Phi_{\tilde{\epsilon}} = c/\bar{\rho}$ where $\bar{\rho} = 1263$ mg/mL is the ratio of molar mass and molar volume of an isolated NUP98 FG-chain (1-499) in aqueous solution obtained at $\alpha = 0.7$ (*65*) in the Martini model. For each set of ($\tilde{\epsilon}$, $c^{\tilde{\epsilon}}_{\mathrm{dilute}}$, $c^{\tilde{\epsilon}}_{\mathrm{dense}}$) parameters, we used Eq. 20 to determine the corresponding value of $\chi$. Fig. S12 shows the linear relation between the $\tilde{\epsilon}$ and $\chi$. According to the FH model for a chain of $N = 499$ residues, the critical cohesive interaction strength is then $\tilde{\epsilon}_c = 0.417$.

*Molecular dynamics simulations of NUP98 condensates.* With a coarse-grained bead-spring polymer model (*60*) parametrized to match the extension of single NUP98 FG-chains(1-499) and the phase behavior of NUP98 FG-chains, we could sample their large-scale motions and condensate formation. In simulations of 500 chains contained in an elongated box, we found that condensation occurs above a critical interaction strength of $\tilde{\epsilon_c} \approx 0.41$ (Fig. S13A and B). Just below, for $\tilde{\epsilon} = 0.4$, the chains uniformly filled the simulation box. The phase coexistence line in the plane of concentration ($c$) and cohesive strength ($\tilde{\epsilon}$) is captured well by the Flory-Huggins (FH) mean-field model (Fig. S13C). For $\tilde{\epsilon} = 0.44$, the calculated concentrations of the dense and dilute phase ($190.6 \pm 8.7$ mg/mL and $0.98 \pm 0.80$ mg/mL) closely match those measured for NUP98-FG and an engineered 12mer GLFG ($0.3 - 0.5$ mg/mL (*71*, *72*) and 175 mg/mL (*71*), respectively).

The conformations of the chains inside the condensate reproduce the experimental smFRET distance measurements (Fig. S13D). Compared to single chains (Fig. S11), the chains in the condensate are more extended with only a weak dependence on $\tilde{\epsilon}$. The interactions with other chains in effect mimic good-solvent conditions for the individual polymers, akin to the Flory hypothesis for polymer melts. Compared to the chains in the NPC (Fig. 4), the chain distances are nearly insensitive to the interaction strength in the condensate, i.e., for $\tilde{\epsilon} > 0.41$ (Fig. S13D).

75. P.-G. de Gennes, *Scaling concepts in polymer physics* (Cornell university press, 1979).

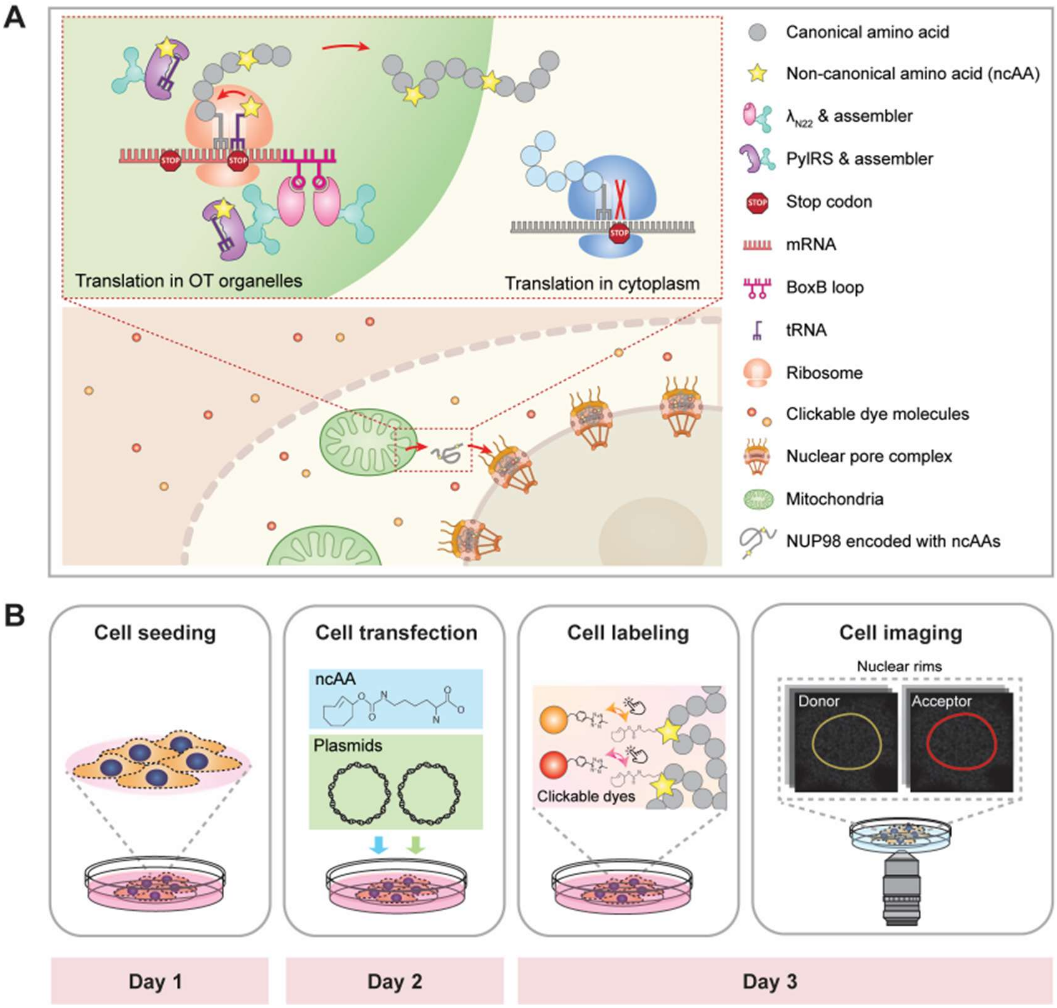


**Fig. S1. Schematics showing the orthogonal translating (OT) organelles enabled genetic code expansion (GCE) system and the sample preparation pipeline. (A)** Schematic of the OT organelles that form a distinct protein translational machinery on the outer mitochondrial membrane surface. The OT organelles exclusively expanded the genetic codon of the target NUP98, ensuring minimal interference with the endogenous protein translation in the cytoplasm. Non-canonical amino acids (ncAAs) were introduced into NUP98 at sites, specified by Amber mutation. The NUP98 encoded with ncAAs docked on the nuclear pore complex during mitosis. The ncAAs were labeled with tetrazine-modified organic dyes *via* click chemistry. **(B)** Schematics showing sample preparation pipeline including cell seeding, cell transfection with NUP98 and OT organelle plasmids, labeling the incorporated ncAAs with clickable dyes, and fluorescent lifetime imaging of labeled NUP98 on the nuclear rims using the custom-built FLIM-FRET optical setup.

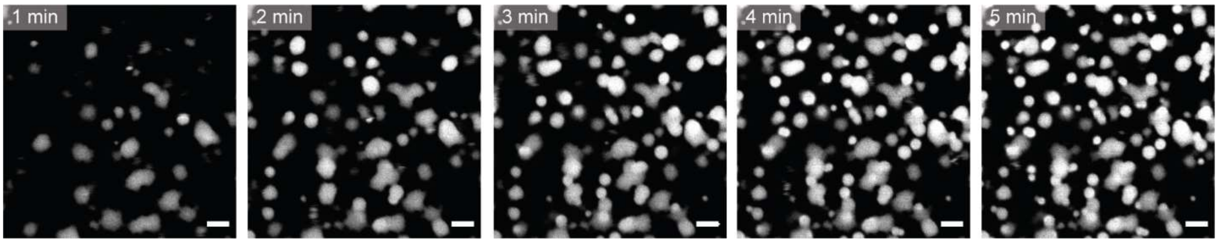


**Fig. S2. Time-lapse images showing phase separation of the purified NUP98 FG-domain *in vitro*.** The early stage of the formed condensates demonstrated liquid-like behavior where fast merging events were observed. Scale bar 5 µm.

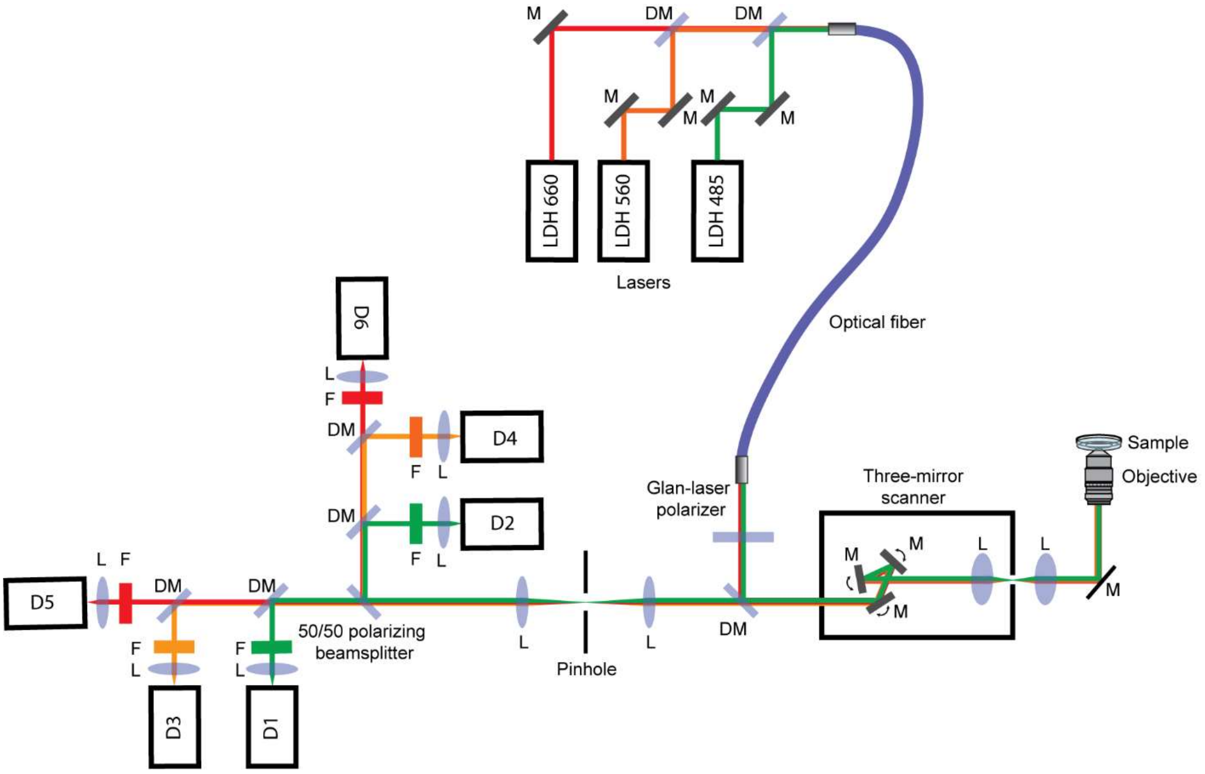


**Fig. S3. Schematic of the custom-built FLIM-FRET imaging system.** Picosecond pulsed lasers diode heads including the wavelengths of 485 nm, 560 nm, and 660 nm were controlled through a multichannel picosecond diode laser driver. The beams were coupled into a single-mode polarization-maintaining optical fiber. The beam traveled through a Glan-laser polarizer and was directed into a laser scanning system. The three galvo mirrors in the scanning system were imaged onto the back focal plane of the objective with a 200 mm tube lens. The fluorescence emission was focused onto a pinhole, and then separated into parallel and perpendicular components using a 50/50 polarizing beam splitter. Each component was further separated by two sets of beamsplitters, passed through three sets of bandpass filters, and focused onto the single-photon counting detectors. The signals from the photon detectors were recorded by a TCSPC system. LDH, picosecond laser diode heads; L, lens; M, mirror; DM, dichroic mirror; D1, D3, D5, detectors perpendicular to the laser excitation; D2, D4, D6, detectors parallel to the laser excitation.

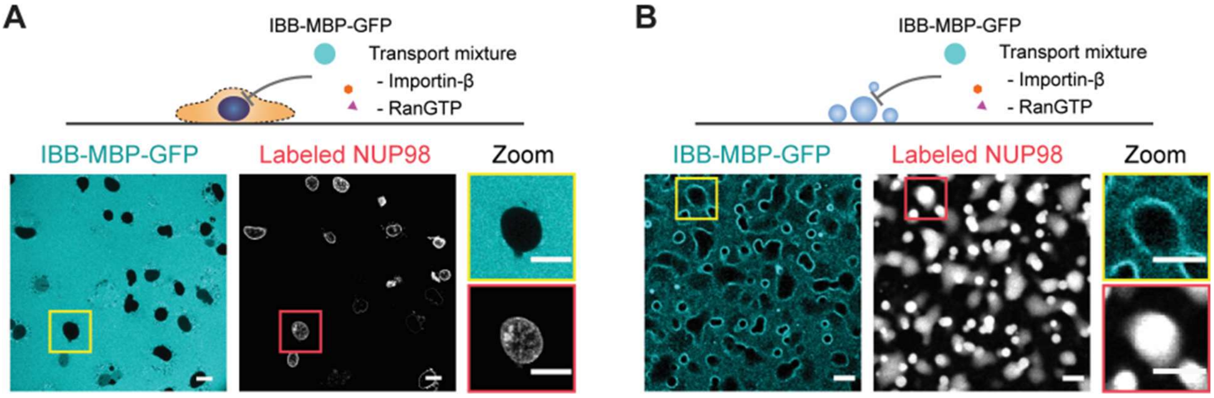


**Fig. S4. Control experiments of transport assay.** IBB-MPB-GFP cargo in the transport mixture without Importin-β and RanGTP was not imported into **(A)** the labeled COS-7 cells or **(B)** the *in vitro* NUP98 FG-condensates.

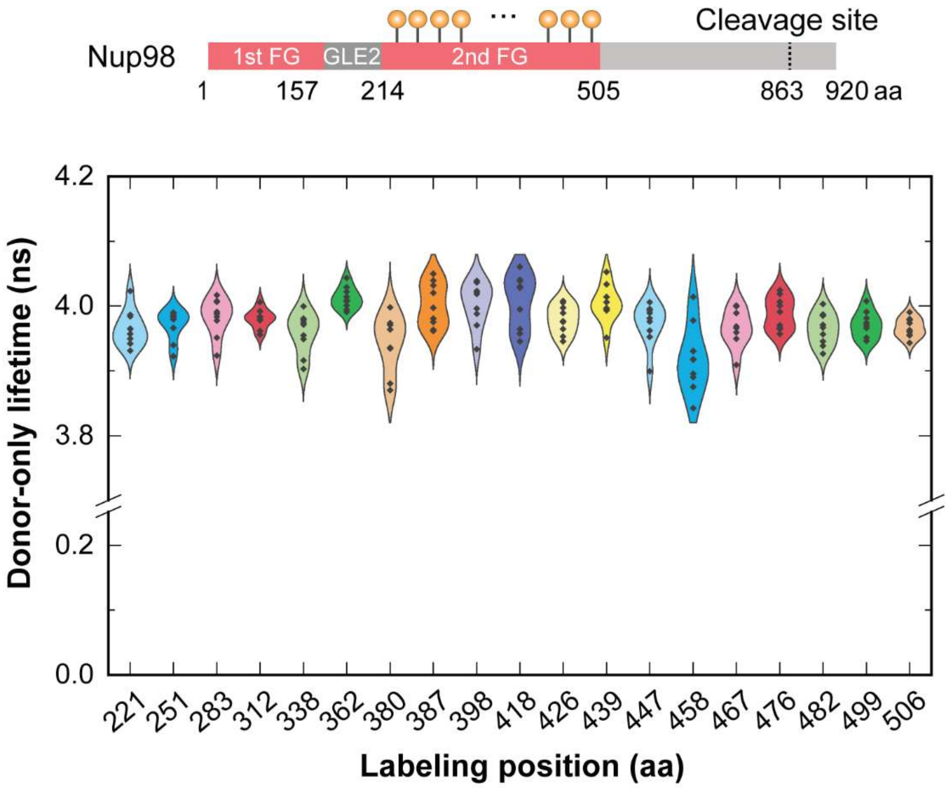


**Fig. S5. Measurements of donor-only lifetime among different labeled sites inside the NPC.** Different positions along the FG-domain of NUP98 labeled with the donor dye showed practically the same lifetime inside the NPC, indicating the fluorophores at different labeling sites shared the same fluorescent properties.

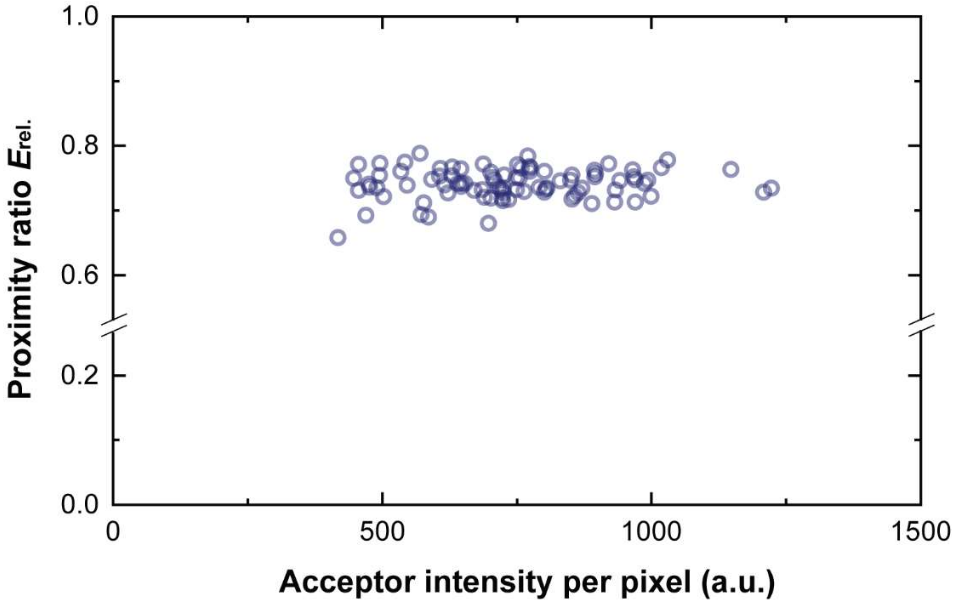


**Fig. S6. The relative FRET efficiency *versus* the acceptor intensity per pixel (excited by 660 nm laser) on the nuclear rim for double-labeled mutant NUP98$^{A221TAG\text{-}I439TAG}$.** To ensure that cells with similar expression levels and not highly overexpressed mutant NUP98 were chosen (which typically leads to large visible aggregates), we always checked the acceptor intensity per pixel (excited by 660 nm laser) to estimate the expression level of the mutant NUP98 before we started the FRET measurements. Within the selected intensity range, on the nuclear rim, the relative FRET efficiency did not corelate with the acceptor intensity per pixel excited by 660 nm laser, indicating that no intermolecular FRET was measured.

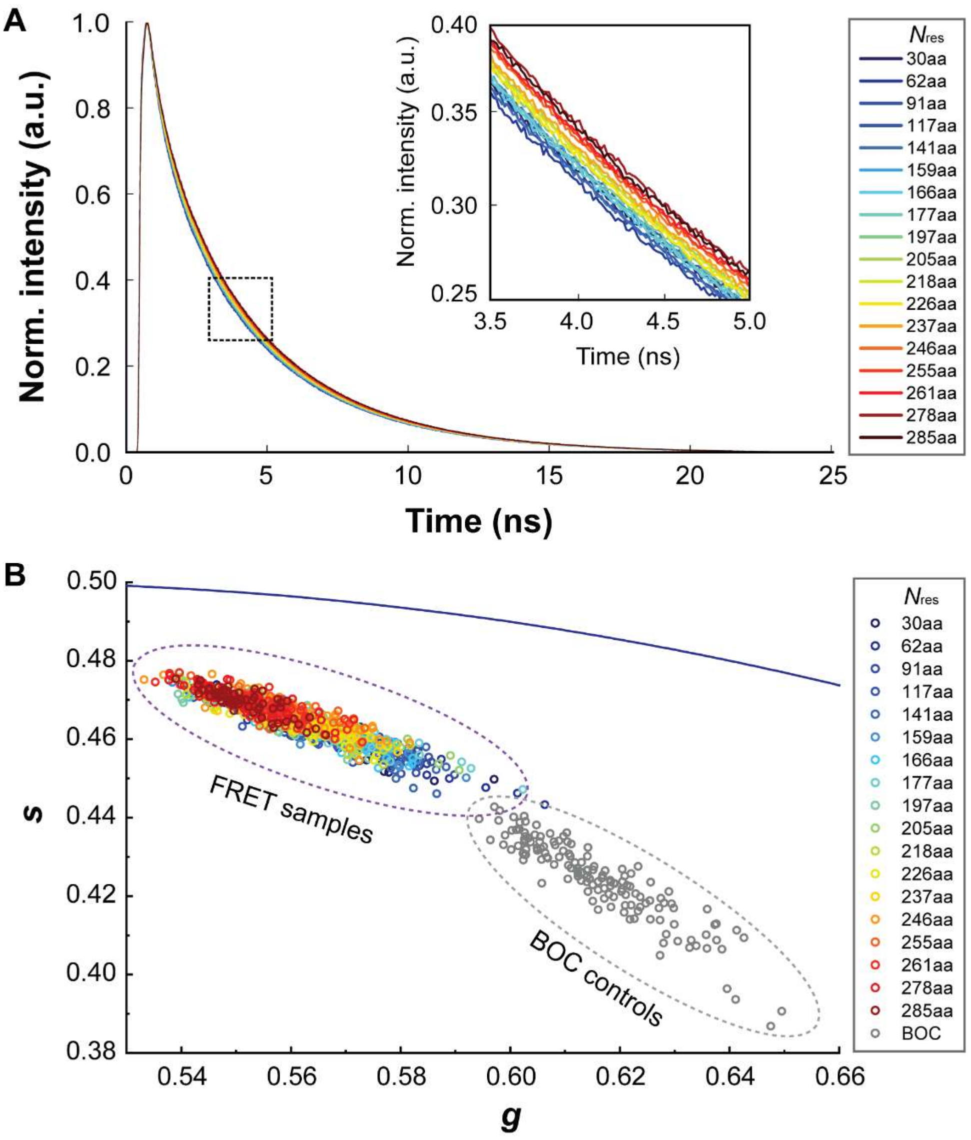


**Fig. S7. Fluorescent lifetime analysis of the eighteen FLIM-FRET probed chain segments of NUP98 FG-domain inside the NPC.** (A) Normalized donor fluorescence intensity profiles of the eighteen chain segments of NUP98 FG-domain for the FLIM-FRET measurements. Each profile represents an averaged result of ~100 cells. (B) Extended phasor plot for Fig. 2G where the lifetime decays of NUP98 encoded with a nonreactive ncAA Lys-BOC are also included. The lifetime decays of the FRET samples could be well separated from the BOC controls. The averaged lifetime decay of the BOC controls was used to determine the cellular background signal for the lifetime fitting procedure.

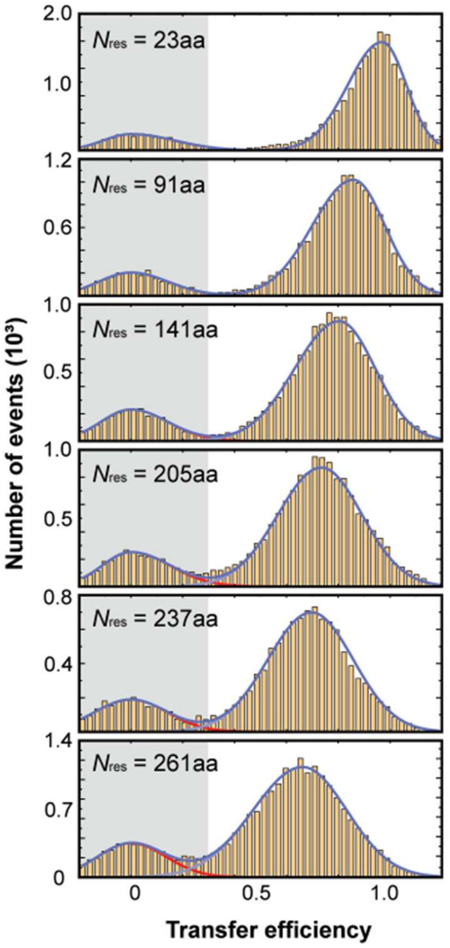


**Fig. S8. Single-molecule FRET measurements in solution**. FRET efficiency histograms for different mutants of NUP98 FG-domain in 1x PBS. The peaks in the grey shadow indicate the donor-only population. The FRET population was fitted with a bi-Gaussian function to retrieve the FRET efficiency.

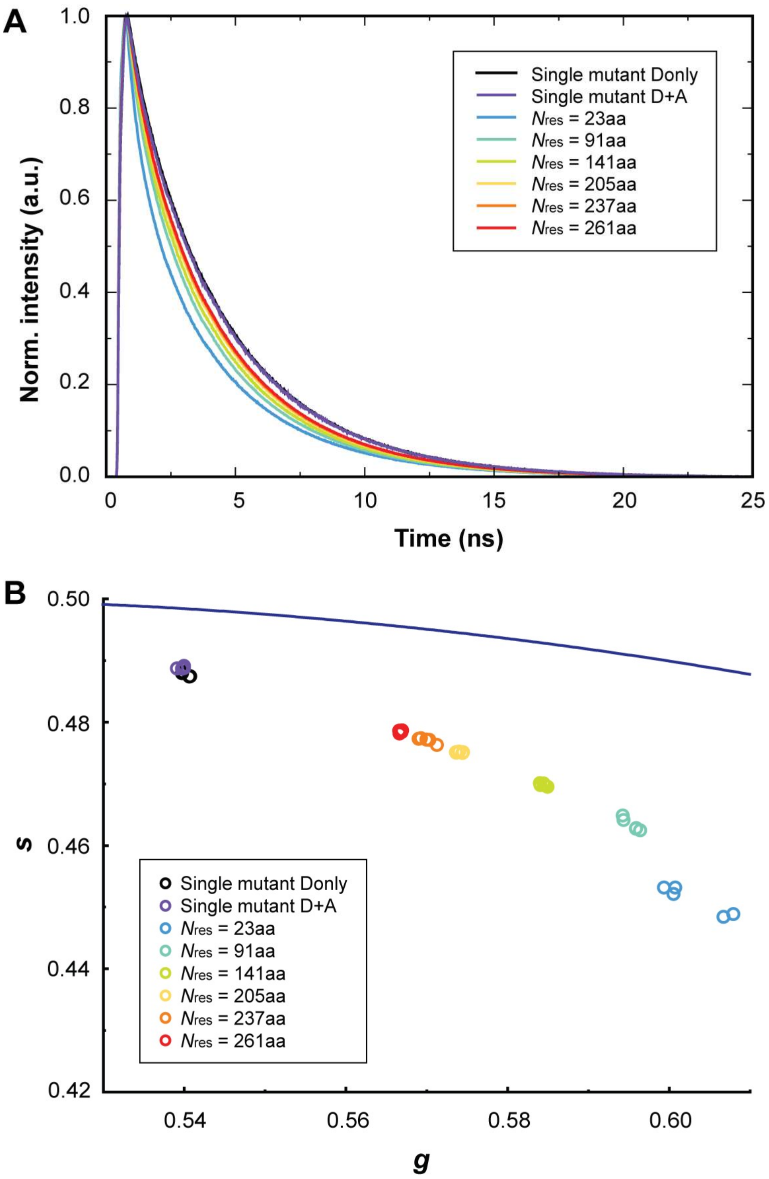


**Fig. S9. FLIM-FRET analysis of NUP98 FG-condensates *in vitro*. (A)** Normalized donor fluorescence intensity profiles of six NUP98 double-mutants in phase-separated condensates that were allowed to settle on the coverslip ($N = 5$). $N_{\mathrm{res}}$ refers to the number of amino acid residues between the two labeled sites. The normalized donor fluorescence intensity profiles of the single-mutant NUP98$^{\mathrm{A221C}}$ labeled with donor only or donor and acceptor mixture showed no difference, indicating no inter-molecular FRET was detected and the lifetime differences detected in the double-mutants were due to intra-molecular FRET. **(B)** Phasor plot showing the donor lifetimes of the six double-mutants for individual experimental repeats ($N = 5$). On the same phasor plot, the lifetimes for the single-mutant labeled with donor-only and a mixture of donor and acceptor were also compared, which showed no difference and ensured no inter-molecular FRET was involved during the measurements.

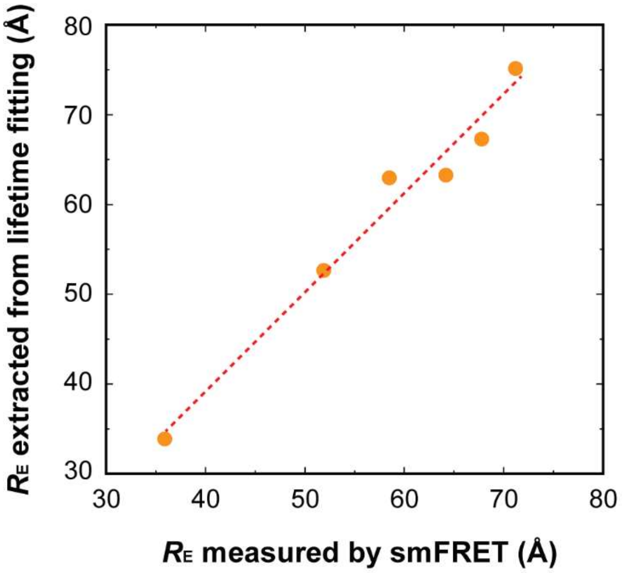


**Fig. S10. Correlation between the end-to-end distance $R_E$ measured by intensity-based smFRET and $R_E$ recovered from the lifetime-based fitting pipeline.** In contrast to our in-cell measurements, on the single molecule level FRET efficiency can be measured intensity-based and lifetime-based and from both $R_E$ can be computed independently. From the fluorescence measurements shown in Fig. S8, we fitted with Equation 11 and 12 and extracted the $R_E$ for both strategies. The detected high correlation of $R^2 = 0.97$ validates the use of our FLIM-FRET pipeline to measure $R_E$.

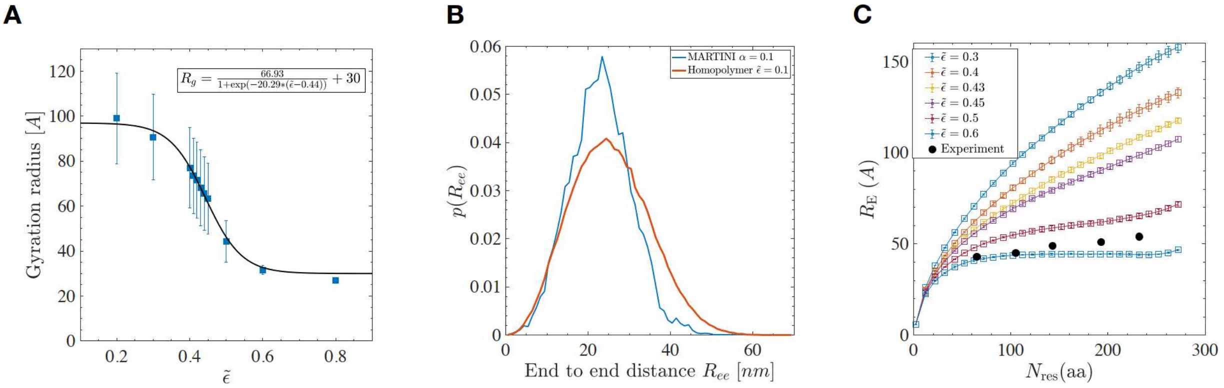


**Fig. S11. Configuration of single NUP98 FG-chain (1-499) from MD simulations. (A)** Radius of gyration of the homopolymer of size $N$ = 499 as a function of cohesive strength $\tilde{\boldsymbol{\epsilon}}$. The solid line shows a logistic function fitted to the data. The error bar shows the standard deviation of the chain's gyration radius calculated over the whole simulation time ($\mathbf{1.5 \times 10^6 \tau}$). The coil-globule transition occurs at $\tilde{\boldsymbol{\epsilon}} = \mathbf{0.44}$. **(B)** Probability distribution of the end-to-end distance for the homopolymer model ($\tilde{\boldsymbol{\epsilon}} = \mathbf{0.1}$) and Martini model ($\boldsymbol{\alpha} = \mathbf{0.1}$) of NUP98 FG-chain in the limit of weak cohesion. **(C)** Mean distance in single NUP98 FG-chain (1-499) from MD simulations versus residue separation. Solid symbols show the results of the smFRET measurements.

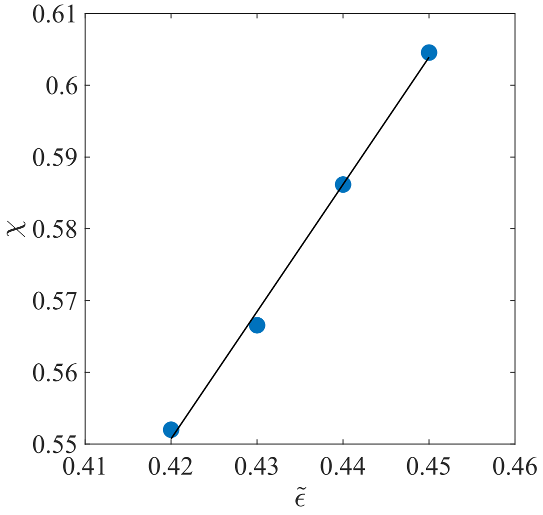


**Fig. S12. Linear relation between the Flory parameter $\chi$ from Equation 20 and the cohesive strength $\tilde{\epsilon}$ of NUP98-FG (1-499).** The line shows a linear fit with coefficients slope $A$ = 1.773 and intercept $B$ = -0.1938.

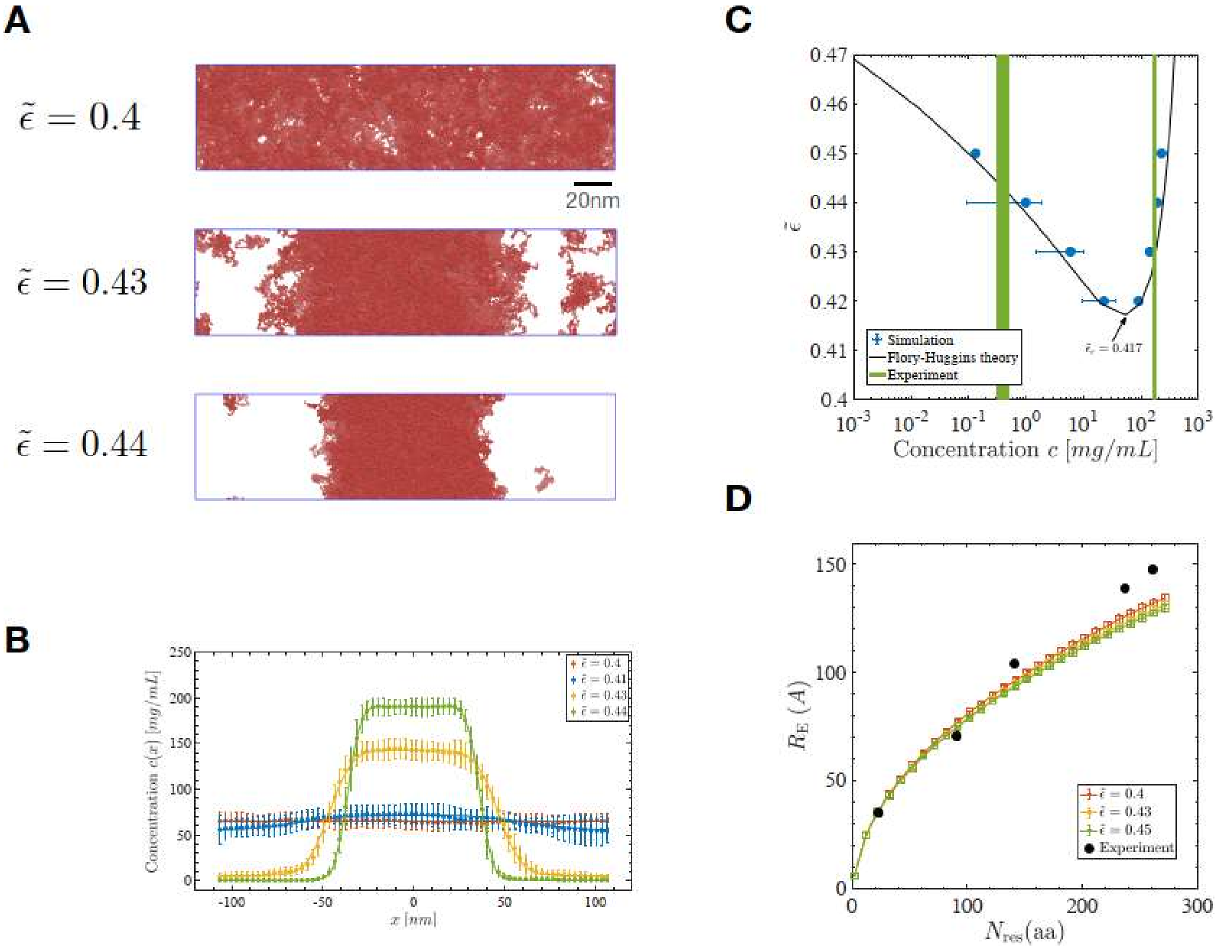


**Fig. S13. Coarse-grained MD simulations of NUP98 FG-condensates. (A)** Condensate formation of homopolymer model of NUP98-FG at different cohesive interaction strengths $\tilde{\boldsymbol{\epsilon}}$. Shown are side-views of elongated simulation box (blue outline) with 500 NUP98-FG (1-499) chains (red). **(B)** Concentration profiles of the NUP98-FG system at different cohesive interaction strengths. For $\tilde{\boldsymbol{\epsilon}} = \mathbf{0.4}$, just below the critical interaction strength, the chains are uniformly distributed inside the system. **(C)** Phase diagram of the NUP98 FG-condensate in the plane of cohesive interaction strength $\tilde{\boldsymbol{\epsilon}}$ and the concentration $\boldsymbol{c}$ of coexisting phases. The solid black line is the result of Flory-Huggins theory for a homopolymer of a length $N = 499$. The critical cohesive strength is obtained as $\tilde{\boldsymbol{\epsilon}}_{\boldsymbol{c}} \approx \mathbf{0.41}$. The experimental values of the dense and dilute phases are marked in green bars (*71*, *72*). **(D)** Mean distance $\boldsymbol{R}_{\mathbf{E}}$ of beads on the same NUP98-FG chain inside the condensate as function of residue separation $\boldsymbol{N}_{\mathbf{res}}$.

**Table S1. Plasmids used in this work**

| Construct | In Figure |
|---|---|
| pcDNA3.1-FLAG-hsNUP98$^{A221TAG}$-boxB | Fig.1-3, Fig.S4 |
| pcDNA3.1-FLAG-hsNUP98$^{A221TAG-A251TAG}$-boxB | Fig.2-3, Fig.S7 |
| pcDNA3.1-FLAG-hsNUP98$^{A221TAG-S283TAG}$-boxB | Fig.2-3, Fig.S7 |
| pcDNA3.1-FLAG-hsNUP98$^{A221TAG-S312TAG}$-boxB | Fig.2-3, Fig.S7 |
| pcDNA3.1-FLAG-hsNUP98$^{A221TAG-S338TAG}$-boxB | Fig.2-3, Fig.S7 |
| pcDNA3.1-FLAG-hsNUP98$^{A221TAG-S362TAG}$-boxB | Fig.2-3, Fig.S7 |
| pcDNA3.1-FLAG-hsNUP98$^{A221TAG-A380TAG}$-boxB | Fig.2-3, Fig.S7 |
| pcDNA3.1-FLAG-hsNUP98$^{A221TAG-S387TAG}$-boxB | Fig.2-3, Fig.S7 |
| pcDNA3.1-FLAG-hsNUP98$^{A221TAG-S398TAG}$-boxB | Fig.2-3, Fig.S7 |
| pcDNA3.1-FLAG-hsNUP98$^{A221TAG-A418TAG}$-boxB | Fig.2-3, Fig.S7 |
| pcDNA3.1-FLAG-hsNUP98$^{A221TAG-A426TAG}$-boxB | Fig.2-3, Fig.S7 |
| pcDNA3.1-FLAG-hsNUP98$^{A221TAG-I439TAG}$-boxB | Fig.2-3, Fig.S7 |
| pcDNA3.1-FLAG-hsNUP98$^{A221TAG-A447TAG}$-boxB | Fig.2-3, Fig.S6-7 |
| pcDNA3.1-FLAG-hsNUP98$^{A221TAG-A458TAG}$-boxB | Fig.2-3, Fig.S7 |
| pcDNA3.1-FLAG-hsNUP98$^{A221TAG-A467TAG}$-boxB | Fig.2-3, Fig.S7 |
| pcDNA3.1-FLAG-hsNUP98$^{A221TAG-A476TAG}$-boxB | Fig.2-3, Fig.S7 |
| pcDNA3.1-FLAG-hsNUP98$^{A221TAG-A482TAG}$-boxB | Fig.2-3, Fig.S7 |
| pcDNA3.1-FLAG-hsNUP98$^{A221TAG-S499TAG}$-boxB | Fig.2-3, Fig.S7 |
| pcDNA3.1-FLAG-hsNUP98$^{A221TAG-M506TAG}$-boxB | Fig.2-3, Fig.S7 |
| pcDNA3.1-TOM20$_{1-70}$-FUS$_{1-478}$-4x$\lambda_{N22}$-PylRS$^{Y306A,Y384F}$_U6-tRNA$^{Pyl}$ | Fig.1-3, Fig.S4-7 |
| pcDNA3.1-FLAG-hsNUP98$^{A251TAG}$-boxB | Fig.S5 |
| pcDNA3.1-FLAG-hsNUP98$^{S283TAG}$-boxB | Fig.S5 |
| pcDNA3.1-FLAG-hsNUP98$^{S312TAG}$-boxB | Fig.S5 |
| pcDNA3.1-FLAG-hsNUP98$^{S338TAG}$-boxB | Fig.S5 |
| pcDNA3.1-FLAG-hsNUP98$^{S362TAG}$-boxB | Fig.S5 |

| | |
|---|---|
| pcDNA3.1-FLAG-hsNUP98$^{A380TAG}$-boxB | Fig.S5 |
| pcDNA3.1-FLAG-hsNUP98$^{S387TAG}$-boxB | Fig.S5 |
| pcDNA3.1-FLAG-hsNUP98$^{S398TAG}$-boxB | Fig.S5 |
| pcDNA3.1-FLAG-hsNUP98$^{A418TAG}$-boxB | Fig.S5 |
| pcDNA3.1-FLAG-hsNUP98$^{A426TAG}$-boxB | Fig.S5 |
| pcDNA3.1-FLAG-hsNUP98$^{I439TAG}$-boxB | Fig.S5 |
| pcDNA3.1-FLAG-hsNUP98$^{A447TAG}$-boxB | Fig.S5 |
| pcDNA3.1-FLAG-hsNUP98$^{A458TAG}$-boxB | Fig.S5 |
| pcDNA3.1-FLAG-hsNUP98$^{A467TAG}$-boxB | Fig.S5 |
| pcDNA3.1-FLAG-hsNUP98$^{A476TAG}$-boxB | Fig.S5 |
| pcDNA3.1-FLAG-hsNUP98$^{A482TAG}$-boxB | Fig.S5 |
| pcDNA3.1-FLAG-hsNUP98$^{S499TAG}$-boxB | Fig.S5 |
| pcDNA3.1-FLAG-hsNUP98$^{M506TAG}$-boxB | Fig.S5 |
| pQE-14His-TEV-hsNUP98$_{FG\ 1\text{-}505,\ \Delta GLEBS}$ | Fig.1, Fig.S2, Fig.S4, Fig.S8 |
| pQE-14His-TEV-hsNUP98$_{FG\ 1\text{-}505,\ \Delta GLEBS}^{A221C}$ | Fig.1, Fig.S2, Fig.S4, Fig.S8 |
| pQE-14His-TEV-hsNUP98$_{FG\ 1\text{-}505,\ \Delta GLEBS}^{A221C\text{-}S244C}$ | Fig.1, Fig.S2, Fig.S4, Fig.S8-10 |
| pQE-14His-TEV-hsNUP98$_{FG\ 1\text{-}505,\ \Delta GLEBS}^{A221C\text{-}S312C}$ | Fig.1, Fig.S2, Fig.S4, Fig.S8-10 |
| pQE-14His-TEV-hsNUP98$_{FG\ 1\text{-}505,\ \Delta GLEBS}^{A221C\text{-}S362C}$ | Fig.1, Fig.S2, Fig.S4, Fig.S8-10 |
| pQE-14His-TEV-hsNUP98$_{FG\ 1\text{-}505,\ \Delta GLEBS}^{A221C\text{-}A426C}$ | Fig.1, Fig.S2, Fig.S4, Fig.S8-10 |
| pQE-14His-TEV-hsNUP98$_{FG\ 1\text{-}505,\ \Delta GLEBS}^{A221C\text{-}A458C}$ | Fig.1, Fig.S2, Fig.S4, Fig.S8-10 |
| pQE-14His-TEV-hsNUP98$_{FG\ 1\text{-}505,\ \Delta GLEBS}^{A221C\text{-}A482C}$ | Fig.1, Fig.S2, Fig.S4, Fig.S8-10 |

**Table S2. List of the FG-NUPs and their grafting sites.** Dynamic and frozen regions have type numbers >1 and =1, respectively. The coordinates of all NUPs with "mol" and "type" numbers are provided in a supplementary file.

| Name of NUP | Number of copies | Beginning/ ending residues | Type number of dynamic region in coordinate file | Type number of frozen region in coordinate file | Grafting residues of dynamic region(s) | Frozen residue number range |
|---|---|---|---|---|---|---|
| NUP54 | 32 | 2-493 | 2 | 1 | PRO111 | 111-493 |
| NUP58 | 32 | 2-599 | 3 | 1 | ASN246, LEU418 | 246-418 |
| NUP62 | 40 | 2-502 | 4 | 1 | ALA331 | 331-502 |
| NUP98 | 48 | 2-615 | 6 | 1 | LYS595 | 595-615 |
| NUP214 | 8 | 700-2090 | 5 | 1 | SER974 | 700-974 |
| NUP358 | 40 | 4-3224 | 9 | 1 | SER757 | 1-757 |
| Scaffold NUPs | - | - | - | 1 | - | all |

*Not included in the model are NUP153 and Pom121.

**Movie S1.** MD simulation showing that NUP98 FG-domain phase-separates and forms condensates when $\tilde{\boldsymbol{\epsilon}} = \mathbf{0.44}$.

**Movie S2.** MD simulation of entire Nuclear Pore Complexes. Upper panel= top view, bottom panel = side view. The scaffold structure known from cryo-ET is rendered in blue. The simulations show that FG-NUPs (red) are too loose to generate a polymer network in the NPC when $\tilde{\boldsymbol{\epsilon}} = \mathbf{0.35}$.

**Movie S3.** MD simulation of entire Nuclear Pore Complexes. Upper panel = top view, bottom panel = side view. The scaffold structure known from cryo-ET is rendered in blue. The simulations show that FG-NUPs (red) form extended coil configurations and the inner-ring FG-NUPs fluctuate extensively to form a dynamic barrier across the central channel in the NPC when $\tilde{\boldsymbol{\epsilon}} = \mathbf{0.4}$

**Movie S4.** MD simulation of entire Nuclear Pore Complexes. Upper panel = top view, bottom panel = side view. The scaffold structure known from cryo-ET is rendered in blue. The simulations show that FG-NUPs (red) form a surface condensate and collapse onto the scaffold in the NPC when $\tilde{\boldsymbol{\epsilon}} = \mathbf{0.44}$

**Movie S5.** MD simulation showing that a single NUP98 FG-chain adopts a globular-like structure in aqueous solution when $\tilde{\boldsymbol{\epsilon}} = \mathbf{0.6}$.
